# CD81 is an Ebola virus inhibiting factor that is antagonized by GP and VP40

**DOI:** 10.64898/2026.02.04.703765

**Authors:** Dan Hu, Elena Hagelauer, Lisa Wendt, Matteo Bosso, Maximilian Bunz, Julia Kammerloher, Jasmin S. Kutter, Johannes Brandi, Lina Widerspick, Thomas Hoenen, Michael Schindler

## Abstract

Viruses manipulate the host cell membrane of infected cells for evasion of antiviral immunity, prevention of superinfection and optimization of viral replication and spread. The Ebola virus glycoprotein (EBOV GP) mediates virus entry, but is also known as important factor for subversion of the host’s antiviral immune response. We characterized the dysregulation of cell surface-residing proteins by EBOV GP and found that among several membrane proteins GP interferes with the tetraspanins CD81, CD63 and CD9. This was a conserved function of several filoviral GPs and not observable for viral glycoproteins of other virus families. While CD63 and CD9 were largely dispensable for EBOV replication, CD81 suppressed virus-like particle entry and replication at multiple steps. This phenotype might be explainable by sustained suppression of NFκB by CD81, that is otherwise activated by VP40 and EBOV trVLP replication. We further demonstrate that not only GP but also VP40 interferes with CD81 functionality and that antibody-mediated clustering of CD81 suppresses EBOV infection. Altogether, the tetraspanin CD81 emerges as druggable NFκB and EBOV-inhibiting factor, supporting an important role of NFκB in EBOV replication and potentially virus-induced immunopathogenesis.

**Highlights:**

- EBOV GP and VP40 interfere with CD81 cell surface expression
- CD81 suppresses NFκB signaling which is activated by VP40
- CD81 restricts EBOV VLP uptake and replication
- Targeting CD81 by a crosslinking antibody inhibits EBOV infection

## Introduction

Ebola virus (EBOV), a member of the *Filoviridae* family, causes Ebola virus disease in humans with case fatalities from 40% to 60% (1–3). Other members of the *Filoviridae* including Marburg virus (MARV), Sudan virus (SUDV) and Bundibugyo virus (BDBV) are also known to be highly fatal to humans (1, 4). EBOV is filamentous in shape and enveloped with a membrane derived from the host cell plasma membrane (5). A wide range of cell types can be infected by EBOV, such as macrophages, dendritic cells, hepatocytes, endothelial cells and epithelial cells (6).

For cell entry, EBOV can bind to a number of attachment factors via its surface glycoprotein (GP) and phosphatidylserine (PtdSer) which is embedded in the viral envelope (7). Identified attachment factors include the C-type lectins DC-SIGN, L-SIGN, hMGL, LSECtin and MBL (7–11). Furthermore, the PtdSer receptors Tim1, Tim4 and Tyro3 receptor tyrosine kinases (Axl, Dtk, and Mer) can facilitate attachment (7, 12–15). Cell-based assays demonstrate that following attachment, EBOV is internalized mainly through macropinocytosis and transported to NPC intracellular cholesterol transporter 1 (NPC1)-positive endo-lysosomes, where GP is cleaved by cathepsin B/L and subsequently interacts with the main entry receptor NPC1 to induce virus-host membrane fusion and nucleocapsid release (16–21). The EBOV nucleocapsid contains the negative-stranded RNA genome, the RNA polymerase L, the nucleoprotein (NP), VP35, VP30 and VP24 (22–26). The viral genome comprises seven genes encoding seven structural proteins (NP, VP35, VP40, GP, VP30, VP24 and L) and two nonstructural proteins (sGP and ssGP) (27–30). EBOV genome replication and transcription take place in EBOV inclusion bodies (IBs) in the cytoplasm (31–33). IBs were recently characterized as liquid organelles, their formation is dependent on NP oligomerization and they recruit a number of viral as well as host proteins (34–36). Progeny virions are assembled and released from the plasma membrane, which is mainly driven by matrix protein VP40 and enhanced by NP and GP (37, 38).

Tetraspanins are a family of proteins that harbor four transmembrane helices which are connected by three domains, a large and a small extracellular as well as one small intracellular (39, 40). Tetraspanins are known to form tetraspanin-enriched microdomains with other host proteins, including integrins and actin linkers, and they are involved in multiple cellular processes, such as cell adhesion, migration, fusion, protein trafficking and signaling (41–44). They are furthermore canonical components and marker proteins for extracellular vesicles, especially CD81, the related CD9 and CD63 (45). Tetraspanins also play multiple roles in the context of viral infections. CD81 is the main entry receptor of HCV and downregulated upon infection to prevent superinfection of cells and activate pro-survival NFκB signaling (46). In HIV-1 infection, CD81 might serve as assembly hub (47, 48), is enriched in intracellular virus-containing compartments in macrophages (49, 50) and downregulated by HIV-1 Vpu to increase viral particle infectivity (51). Apart from these dual roles in viral infection, in which CD81 might act pro- or antiviral, dependent on the phase of viral replication, CD81 was also described as host-dependency factor, facilitating replication of Chikungunya virus or Herpes simplex virus (52, 53). However not only CD81, but also other tetraspanins as for instance CD151, CD82 or CD63 play multiple roles during the replicative cycle of various viruses (54), which might not come as a surprise given the large functional diversity of tetraspanins in organizing membrane-related processes viruses have to cope with for viral entry, release as well as viral protein translation and maturation.

The viral surface glycoprotein EBOV GP is not only required for viral entry, but also known to modulate host proteins involved in viral pathogenesis and immune evasion (55–61). To identify novel cell surface plasma membrane-residing proteins specifically modulated by EBOV GP, we performed a flow cytometry-based screen. Three tetraspanins CD81, CD63 and CD9 were identified to be targeted by EBOV GP. We here analyzed the role of CD81, CD63 and CD9 in the EBOV life cycle and found CD81 to exert inhibitory activity at multiple steps of EBOV replication.

## Materials and Methods

### Cell culture

HEK293T and HeLa cells were cultured in DMEM with 10% FCS and 1% Penicillin/Streptomycin (P/S). Huh7.5 cells were cultured in DMEM with 10% FCS, 1% non-essential amino acid (NEAA), 1% sodium pyruvate and 1% P/S. CD81KO, CD63KO, CD9KO and the control cells were cultured in medium containing 1 µg/ml puromycin. Monocyte-derived macrophages (MDM) were prepared from PBMCs essentially as described in (62). PBMCs were isolated from blood samples via Ficoll-Paque gradient centrifugation and macrophages were differentiated from PBMCs in macrophage medium (RPMI + 4% human AB serum + 1% NEAA + 1% sodium pyruvate + 1% P/S + 0.4% MEM vitamins). The cells were cultured in an incubator with 5% CO_2_ and 90% relative humidity at 37 °C. MDMs were generated from pseudonymized buffy coats of volunteer blood donors. Blood donors give informed consent for the use of blood and blood-derived products for research use. We do not collect data concerning age and ethnicity, and we comply with all relevant ethical regulations approved by the ethics committee of the University Hospital Tübingen, (IRB no. 507/2017B01 and 612/2020A). All buffy coat donations are received in pseudonymous form and chosen randomly.

### Plasmids and antibodies

Plasmids used in this study are listed in Table 1. EBOV NP, L, VP35, VP30, VP40 and VP24 were ligated into pCG-3*-IRES-eGFP using XbaI and MluI. The plasmids for the trVLP assay were described before (63). EBOV RNA polymerase II driven tetracistronic minigenome plasmid with a GFP reporter is described here (64). Plasmids expressing EBOV GP, VP40, NP and VP35 as well as CD81 fused with the N/C part of Kusabira-green (KG) protein were generated by using BamHI and NotI restriction sites. The antibodies used for flow cytometry (FC) and Western blot (WB) in this project are listed in Table 2.

**Table 1.**
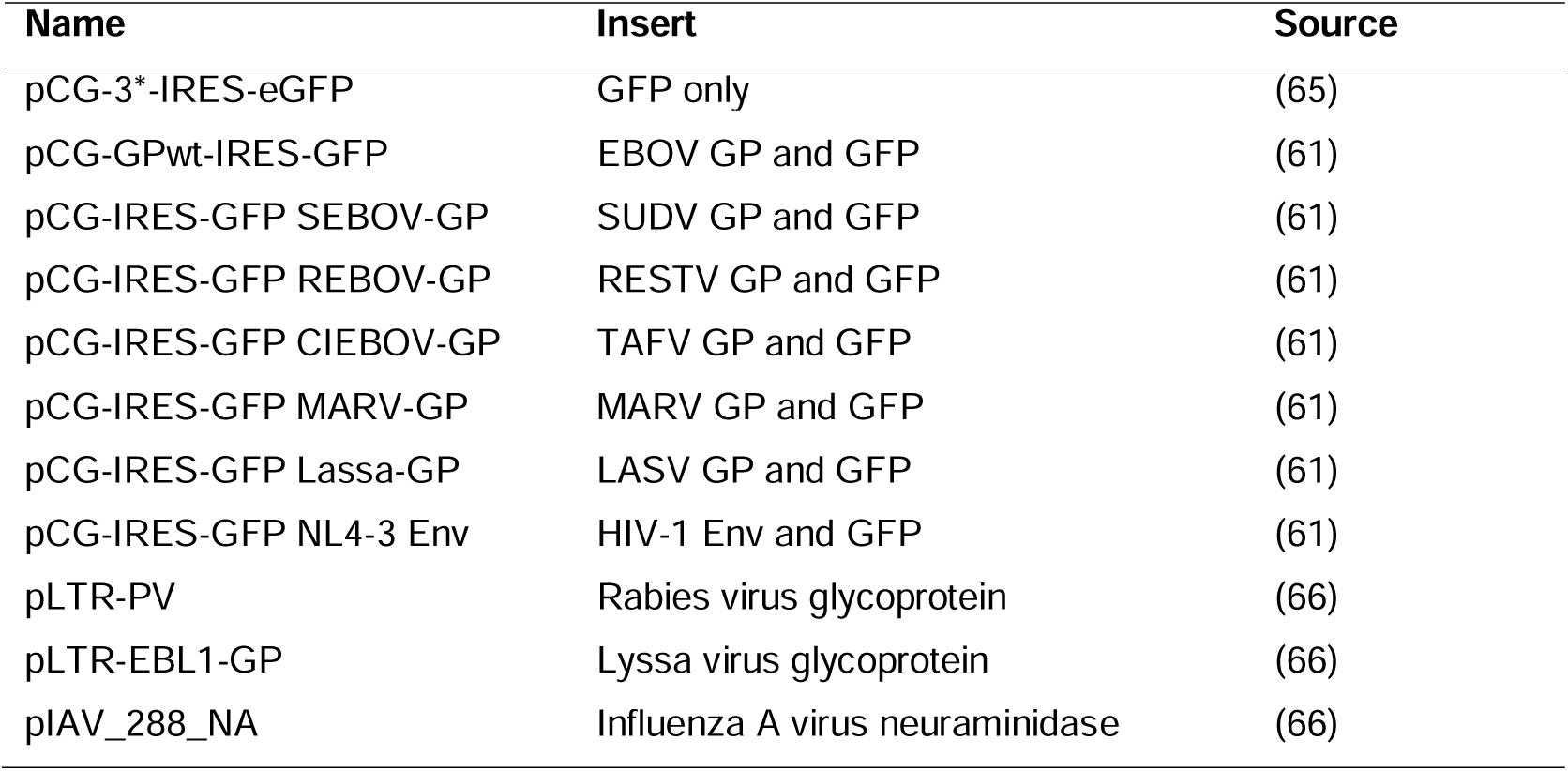
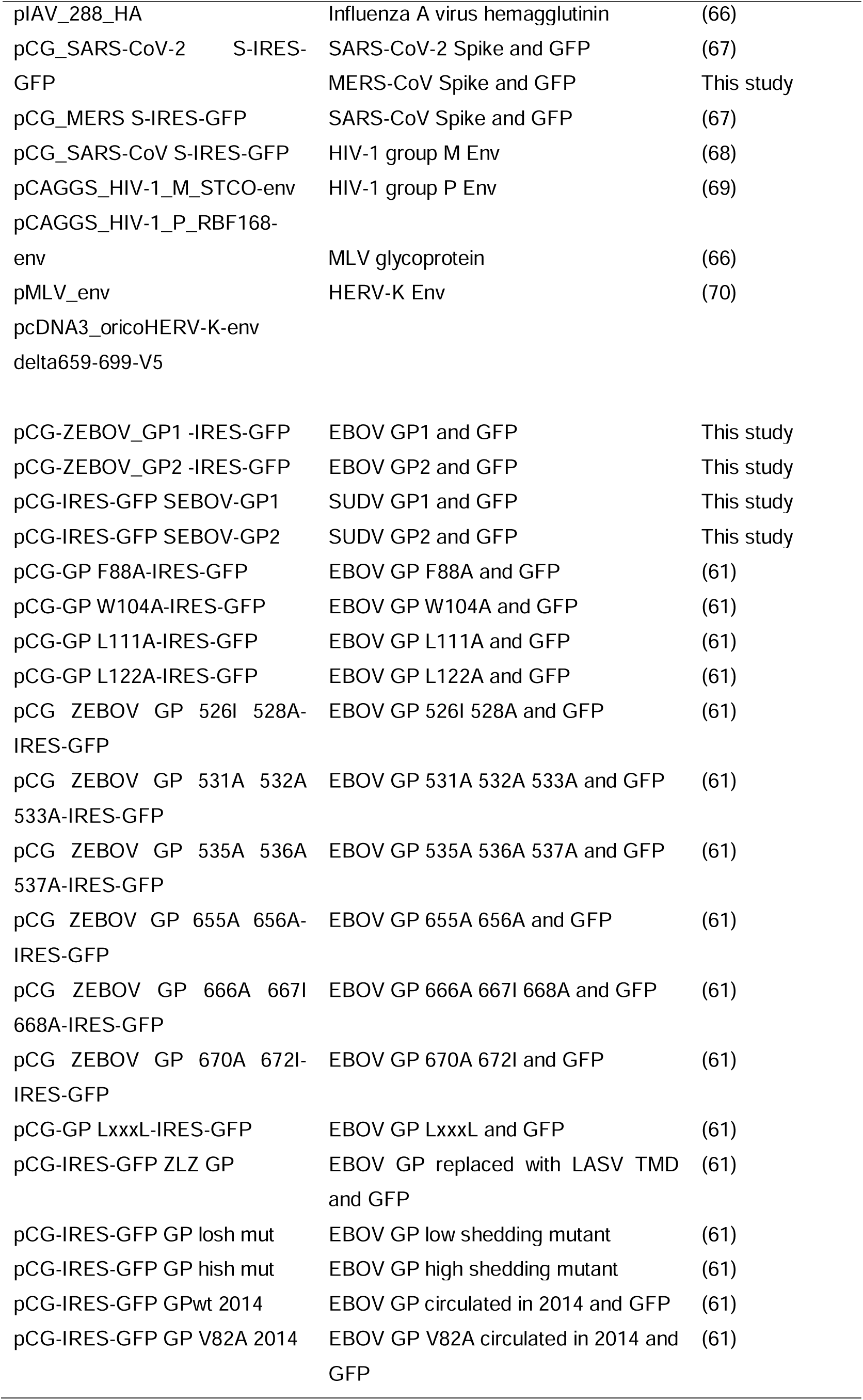
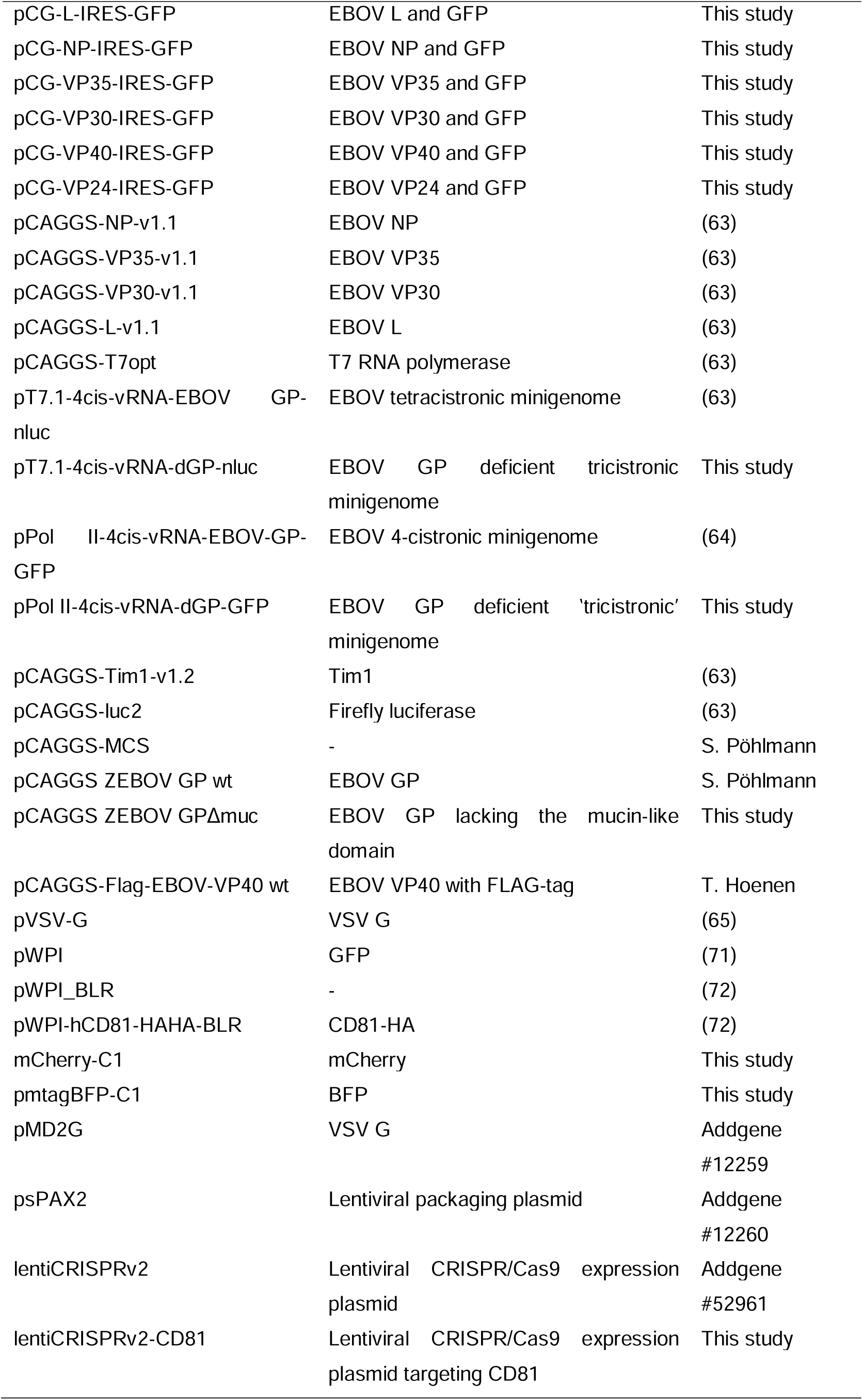
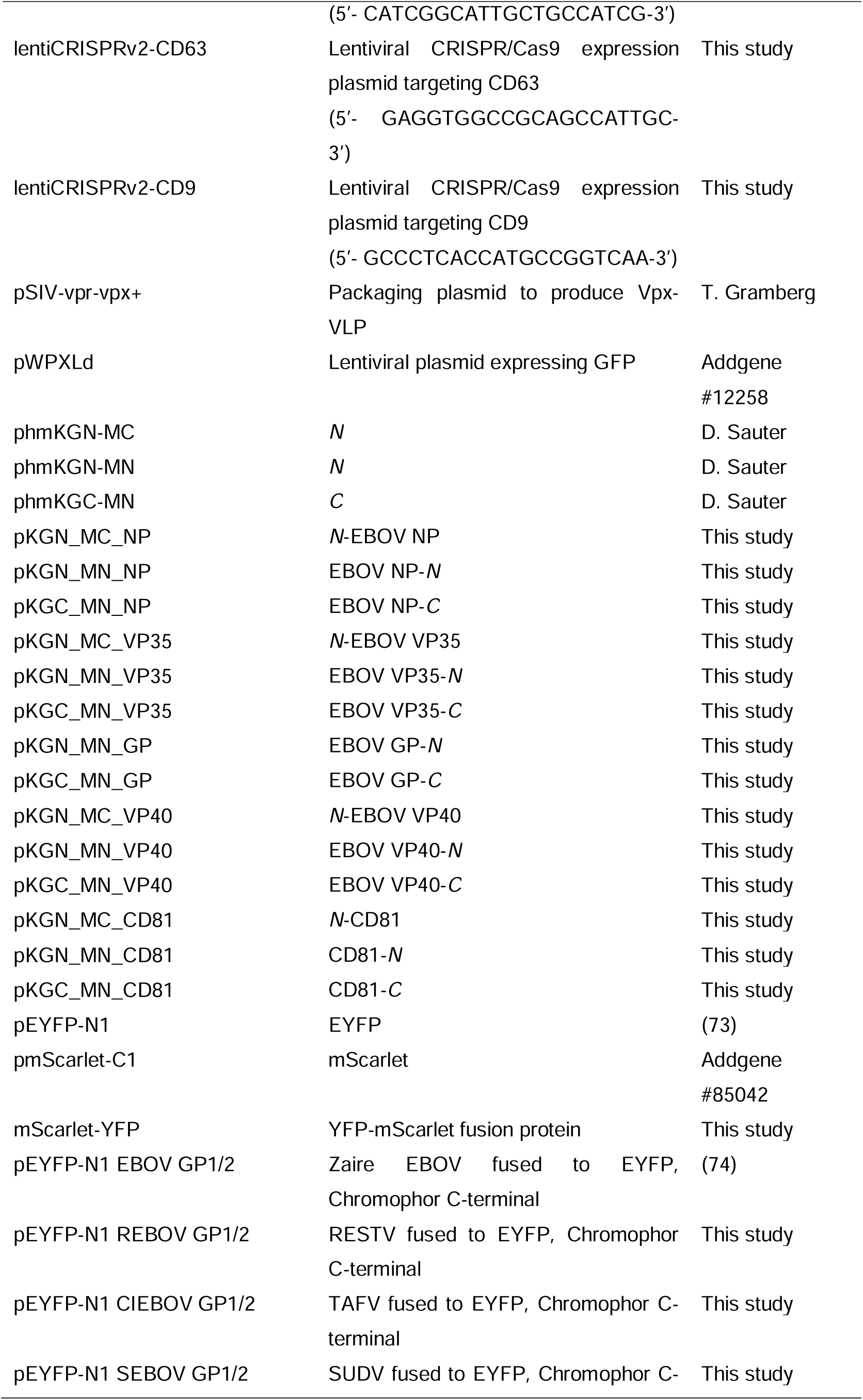
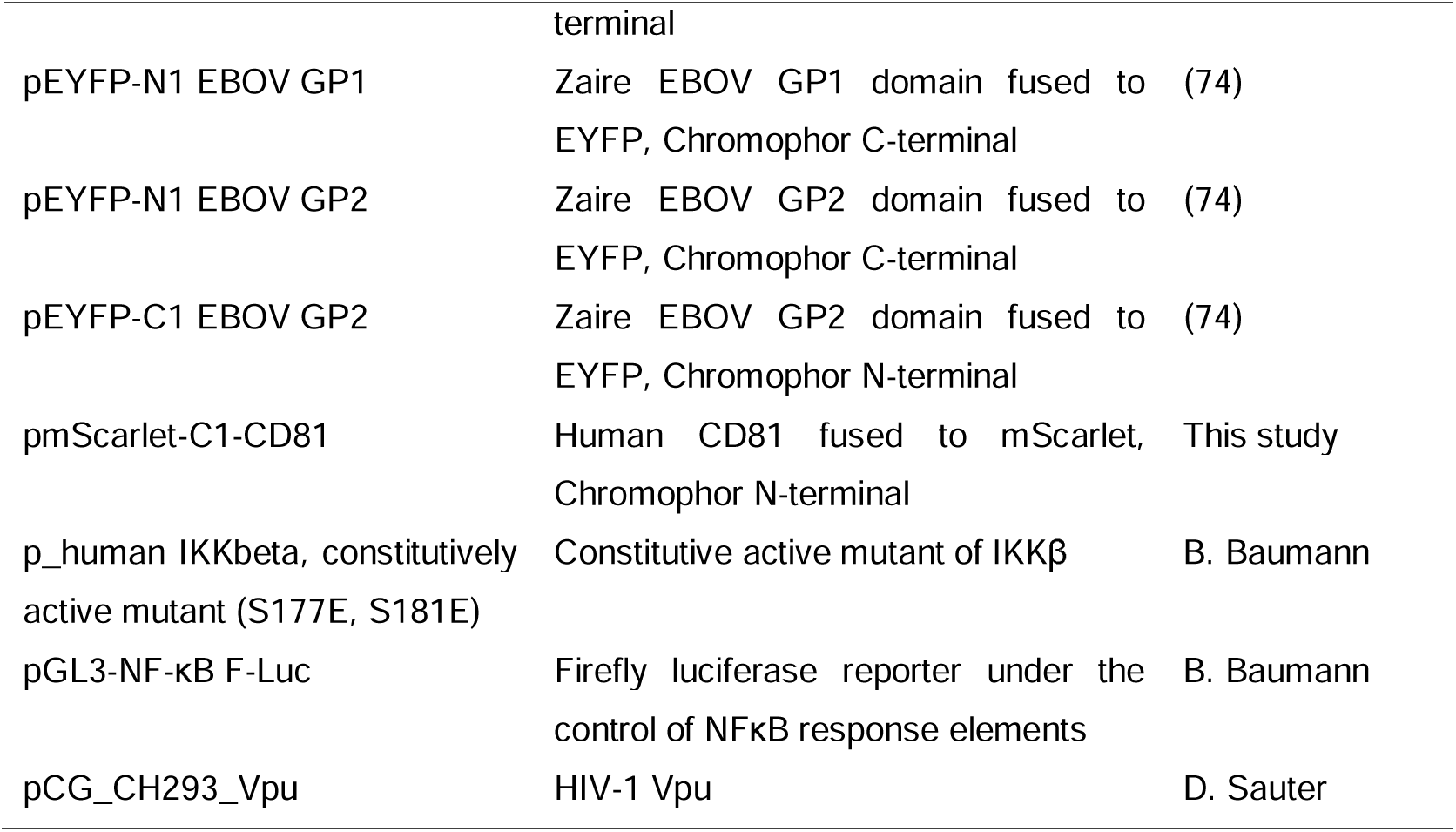
Plasmids.

**Table 2.**
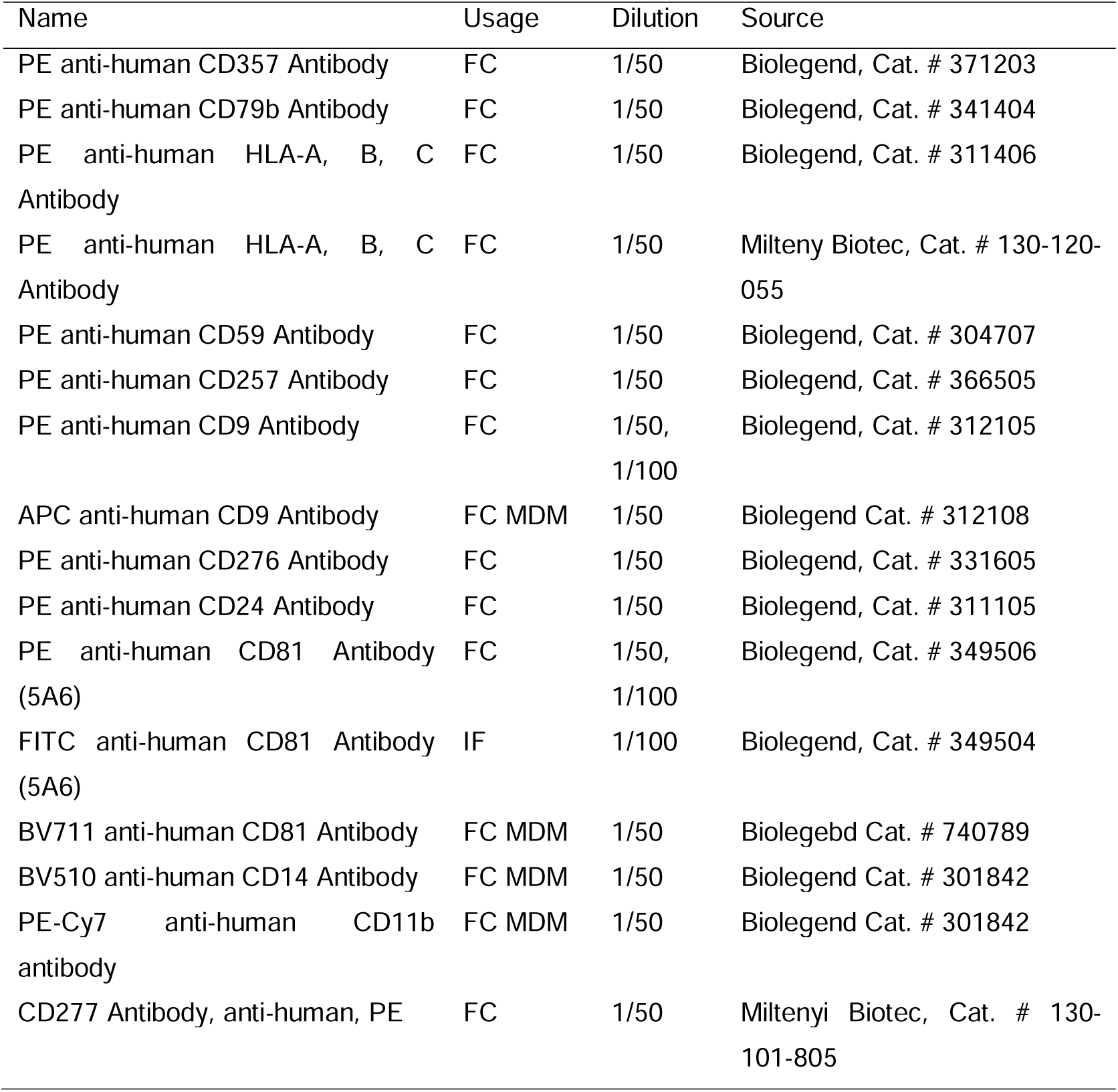
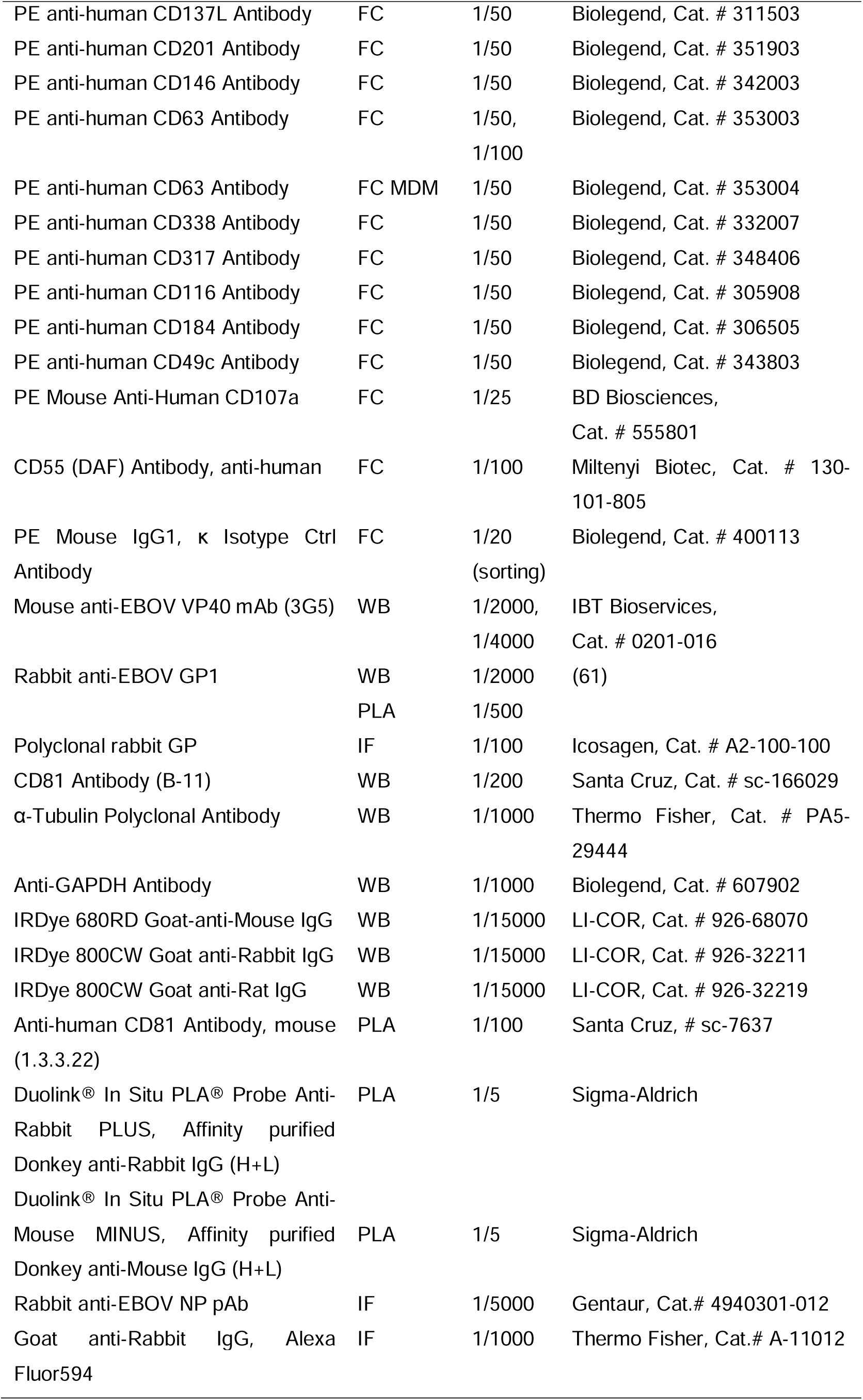
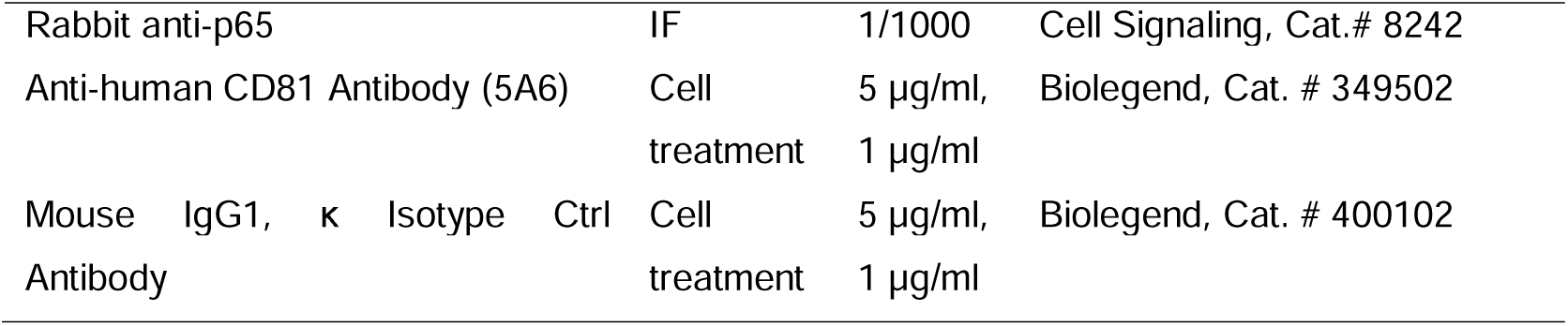
Antibodies.

### Transfection

Calcium phosphate was used for transfection of HEK293T cells. In a 6-well plate format, 5 µg plasmids and 13 µl 2 M CaCl_2_ were mixed with H_2_O in a total volume of 100 µl. The DNA mixture was added dropwise to 100 µl of cold 2× HBS while vortexing. After vortexing for additional 10 s, the transfection mixture was incubated on ice for 15 min and added dropwise to the cells. 6 hours later, the medium was replaced with fresh medium. Polyethylenimine (PEI) was also used for transfection of HEK293T cells. In a 12-well plate format, 1 µg plasmids were mixed with 50 µl Opti-MEM, and 3 µl of 1 µg/µl PEI was mixed with 50 µl Opti-MEM. The PEI mixture was added to the plasmid mixture and mixed by pipetting. After incubation at RT for 15 min, the transfection mixture was added dropwise to the cells. 16 h later (o/n), the medium was replaced with fresh medium. For transfection of other formats, the amount of plasmid and PEI was adjusted accordingly. JetPRIME (PolyPlus) was used for transfection of HEK293T, HeLa and Huh7.5 cells according to the manufacture’s protocol. DharmaFECT 4 transfection reagent was used for transfection of human primary macrophages with ON-TARGETplus human CD81 siRNA – SMARTPool or ON-TARGETplus non-targeting control siRNA #2 according to the manufacturer’s recommended protocol.

### Generation of knockout (KO) cells

HEK293T CD81KO cells were generated via lentiviral transduction. HEK293T cells in 6-well plates were transfected with 0.45 µg pVSV-G, 1.125 µg psPAX2 and 1.5 µg lentiCRISPRv2-CD81 to produce lentiviruses carrying Cas9 and a gRNA targeting CD81. In parallel, HEK293T cells were transfected with 2.5 µg pSIV-vpr-vpx+ and 0.25 µg pVSV-G to produce Vpx-VLPs, which were used to knock down SAMHD1 in target cells to improve lentiviral transduction efficiency (75). 24 h later, the supernatants containing lentiviruses and Vpx-VLPs were collected and centrifuged at 3200 g for 10 min to remove dead cells and cell debris. The cleared supernatants were collected for transduction. HEK293T cells seeded in 6-well plates were treated with Vpx-VLPs for 2 h and infected with lentiviruses for 24 h. 24 h later, the cells were cultured in medium with 1 µg/ml puromycin for selection of successfully transduced cells. HEK293T control, CD63 KO and CD9 KO cells were generated by transfection of HEK293T cells in 6-well plates with 0.45 μg pMD2G, 1.125 μg psPAX2 and 1.5 μg lentiCRISPRv2, lentiCRISPRv2-CD63 or lentiCRISPRv2-CD9. One week later, the cells were cultured in medium containing 1 µg/ml puromycin. After selection, the KO efficiency of the cells was analyzed by flow cytometry. HEK293T CD81 KO cells stained with PE-conjugated anti-human CD81 antibody were subjected for fluorescence-activated cell sorting (FACS), PE mouse IgG1, κ isotype ctrl antibody stained HEK293T CD81 KO cells were used for gating. Collected HEK293T CD81 KO cells from FACS are termed as 293T CD81 KO cells (sorted).

### Legend screen

HeLa cells were transfected with pCG-GPwt-IRES-GFP to coexpress EBOV GP and GFP. 36 hours later, using the LEGENDScreen™ Human Cell PE Kit, the cells were harvested and stained with antibodies against 332 receptors and 10 isotype ctrl antibodies according to the manufacture’s protocol and in detail as previously described (62, 76). The primary data is accessible through sup. Table S1.

### Flow cytometry staining

Surface staining was performed with HEK293T, HeLa, Huh7.5 cells and human primary macrophages. The cells were stained with PE-conjugated antibodies (Table 2) diluted in FACS buffer (1% FCS in PBS) for 30 min at 4 °C in the dark. FACS buffer was added to the cells for washing. The cells were centrifuged at 600 g for 5 min, and the washing step with FACS buffer was repeated once. Subsequently, the cells were either resuspended in FACS buffer for direct measurement or fixed with 2% paraformaldehyde (PFA). For staining of EBOV infected HEK293T cells or MDMs, the cells were fixed as described in EBOV infection before staining. For the PNGase assay, cells were treated 24 h.p.t. for 4 h with 100 U/ml PNGase F (Promega) at 37°C, before harvesting and staining for flow cytometry.

For intracellular staining, 2% PFA fixed (15 min at RT) surface-stained cells were washed with PBS and permeabilized with ice-cold 90% methanol (in H_2_O, stored at -20 °C) at 4 °C for 20 min. PE of surface-stained antibodies is damaged by methanol (77). After the permeabilization, the cells were washed with FACS buffer and blocked with 10% FCS (in PBS) for 30 min at RT. The cells were then stained with PE-conjugated antibodies and washed twice with FACS buffer. For total staining, fixed surface-stained cells were washed with PBS and permeablised with 0.2% saponin for 10 min at RT. The cells were then blocked with 10% FCS in 0.2% saponin for 30 min at RT. After washing with FACS buffer, the cells were stained with PE-conjugated antibodies and washed twice with FASC buffer. For total staining of fixed EBOV infected HEK293T cells, the cells were permeabilized, blocked, stained and washed as described for the intracellular staining.

The measurements were done with the MACSquant VYB (Miltenyi Biotec). For tetraspanin modulation in rgEBOV-GFP-infected MDMs, the 5L Cytek Aurora (Cytek Biosciences) was used. The flow cytometry gating strategy is detailed in sup. Fig. S1 and Fig. 1.

**Fig. 1.**
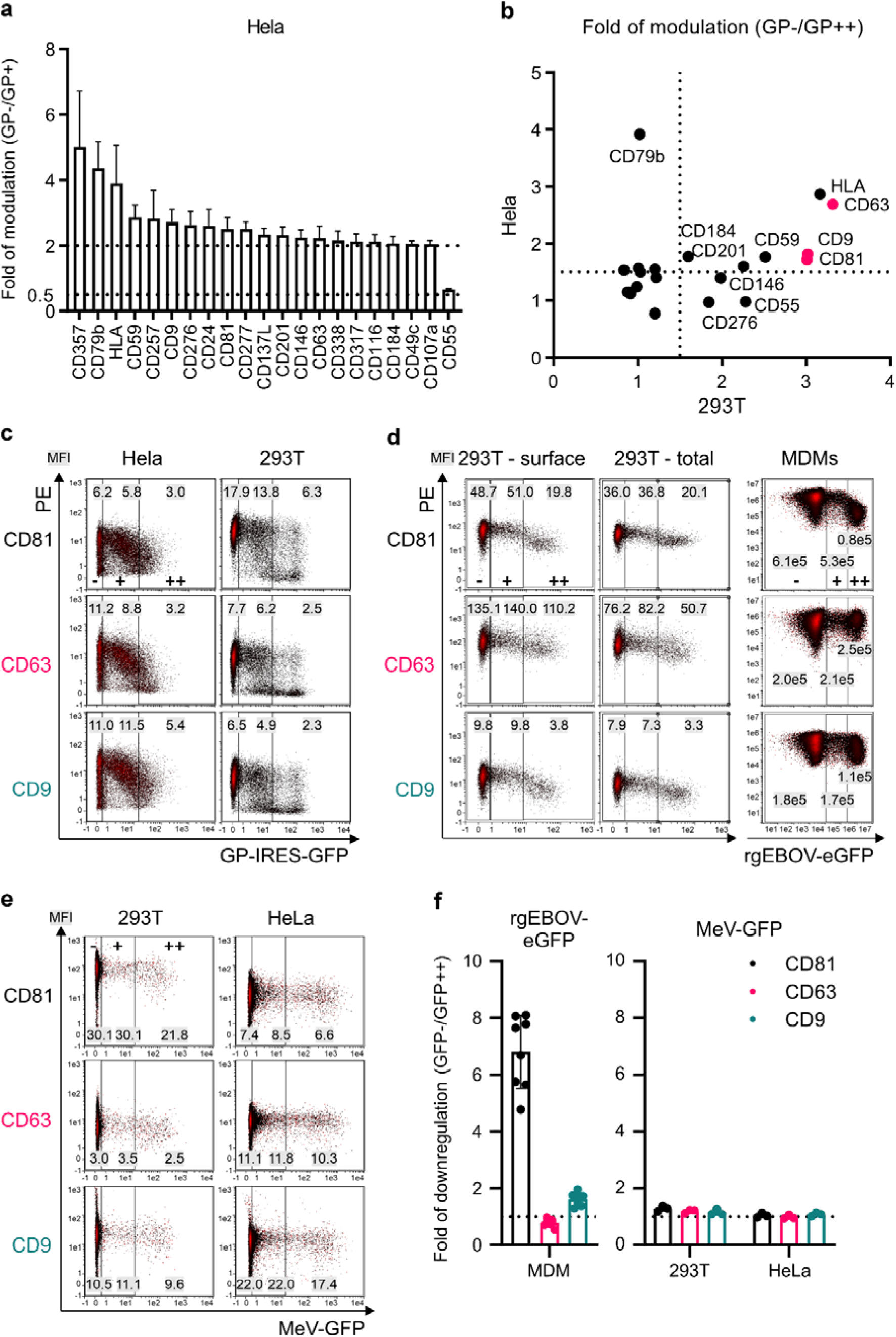
EBOV GP downregulates tetraspanins CD81, CD63 and CD9. (a) EBOV GP modulated surface receptors identified in HeLa cells with a flow cytometry-based screen. Shown are mean values of fold of modulation, calculated by dividing PE MFI of GP negative (GFP-) population by PE MFI of GP positive (GFP+) population, +/- SD (n=3). (b-c) Confirmation of EBOV GP mediated modulation of the identified receptors (b) in 293T and Hela cells with flow cytometry. Shown are (b) mean values of fold of modulation +/- SD (n=3), and (c) representative density plots of cells expressing no (-), medium (+) to high (++) levels of GFP and hence GP (see also Fig. S2a and b). (d, f) 293T cells and monocyte-derived macrophages (MDMs) were infected with authentic rgEBOV-eGFP (MOI=1 and MOI=0.5 respectively). 24 h.p.i. (293T) or 6 d.p.i. (MDMs), the cells were harvested and analyzed for surface and total expression of CD81, CD63 and CD9 by flow cytometry. Shown are (d) representative density plots and (f) mean values of fold of modulation (GFP- /GFP++) +/- SD (n=8 different donors from 2 independent experiments). (e, f) 293T and HeLa cells were infected with Measles virus (MeV-GFP, MOI=0.1). 48 h.p.i., the cells were harvested and analyzed for surface expression of CD81, CD63 and CD9 by flow cytometry. Shown are (e) representative density plots and (f) mean values of fold of modulation (GFP- /GFP++) +/- SD (n=3). Values in the primary FACS plots indicate the PE MFI (mean fluorescence intensity) in the respective region.

### Microscopy and immunofluorescence

For assessment of inclusion body (IB) formation, 1×10^5^ HEK293T control and CD81 KO cells were seeded in poly-L-lysine-coated thin-bottom plates in a 24-well format. The following day, the cells were transfected using PEI with a total of 0.4 µg of DNA. The cells were fixed 24 hours post-transfection (h.p.t.) with 2% PFA for 15 min at RT. After two washing steps with PBS, the cells were permeabilized with 0.2% saponin in PBS for 10 min and subsequently blocked with 10% normal goat serum (NGS) for 1 h at RT. After three washes with PBS, the cells were incubated with the primary antibody for 1 h at RT. Following three additional washes with PBS, the cells were stained with the fluorescently labeled secondary antibody for another hour at RT. Antibody dilutions were prepared in 0.1% saponin in FACS buffer (1% FCS in PBS). To visualize cell nuclei, cells were stained with DAPI (0.1 µg/ml in PBS) for 10 min at RT. After two final washes, images were acquired using the Cytation3 imaging microplate reader at 40x magnification. IBs were quantified as red object count using ImageJ software.

For p65 detection, 2×10^4^ HEK293T control and CD81 KO cells were seeded into a poly-L-lysine-coated 96-well plate and transfected the next day using jetPRIME. 6 h.p.t., the medium was replaced, and the cells were infected with EBOV trVLP-dGP-GFP. 48 h.p.t., the cells were fixed with 2% PFA for 15 min at RT, washed twice with PBS, and incubated for 5 min with a permeabilization and blocking buffer (5% FCS, 0.5% Triton-X100 in PBS). The cells were washed with PBS and incubated with the primary antibody for 1 h at RT. After washing with PBS, the cells were stained with the secondary antibody for 1 h at RT. Nuclei were stained with DAPI (0.1 µg/ml in PBS) for 10 min at RT. The staining procedure concluded with another wash with PBS, and cells were imaged using the Cytation3 at 4x magnification. For quantification using ImageJ, a green mask was generated to identify GFP-positive cells (transfected or infected cells), and the red mean fluorescence intensity (p65 staining) within the green mask was measured.

For assessment of CD81 and GP localization, HeLa cells were seeded in µ-Slide 8 well high glass bottom IBID plates (2×10^4^ cells per well). One day later they were transfected using jetPRIME buffer and medium was changed after 4 h. 24 h.p.t., cells were fixed with 2% PFA, permeabilized with 0.2% saponin and blocked and stained as described before for IB formation detection. Cells were imaged using the ZEISS Apotome 3 fluorescent microscope at 63x magnification.

### Proximity ligation assay (PLA)

HeLa cells were seeded at a density of 1×10^4^ cells/well in a 12-well plate onto cover slips. For transfection, 1.6 µg of plasmid DNA was mixed with 2 µl of Lipofectamine2000, followed by an incubation period of 15 min at RT. Before adding the mixture dropwise to cells, antibiotic free DMEM supplemented with 10% FCS was added to the cells. After an incubation period of 6 h, medium was changed to fresh medium. One day post transfection (d.p.t.), cells were fixed using 2% PFA and subsequently permeabilized with 1% saponin. After blocking cells o/n with 10% FCS, cells were incubated in a humidity chamber with primary antibodies for 1 h at RT. During incubation with primary antibodies, PLA probes were diluted 1:5 in 1% FCS in PBS and incubated for 20 min at RT. After washing cells 2x with wash buffer A, PLA probes were added to the cells and incubated in a humidity chamber at 37 °C for 1 h. Afterwards, cells were again washed 2x with wash buffer A and further incubated with ligation solution in a humidity chamber at 37 °C for 30 min. Subsequently, amplification solution was added to the cells and further incubated at 37 °C for 100 min. Cells were then washed with wash buffer B and dried on a tissue. Cover slips were mounted on microscopic slides, using mounting solution provided with the Duolink kit and analyzed via fluorescence microscopy. Spots were counted using the ImageJ spot counting tool.

### Flow cytometry based-FRET

The experiment was essentially performed as described in (73, 74). HEK293T cells were seeded in a 12-well plate format using 3×10^5^ cells per well and transfected the next day using PEI. For transfection a YFP-fusion protein was co-transfected with a mScarlet-fusion protein and the cells were harvested 24 h.p.t. for flow cytometry analysis. YFP was excited using the 488 nm laser and detected with a 525/50 nm band pass filter. FRET signal was measured by excitation with the 488 nm laser and detection using a 614/50 nm filter. For exclusion of false positive FRET signal, mScarlet was excited with a 561 laser and detected using a 615/20 nm filter.

### Western blot

The cells were lysed with RIPA buffer (140 mM NaCl, 10 mM Tris-HCl, 1 mM EDTA, 0.5 mM EGTA, 0.1% (w/v) Na-Deoxycholate, 0.1% (w/v) SDS, 1% (v/v) TritonX-100 in H_2_O, pH 7.4) with freshly-added protease inhibitor at 4 °C for at least 20 min. The cell lysates were centrifuged at 20000 g for 10 min at 4 °C, and supernatants were taken and mixed with 5× Laemmli buffer. The medium containing trVLPs was cleared by centrifugation at 3200 g for 10 min and the supernatants containing VLPs were taken and top layered on 20% sucrose. The VLPs were pelleted by centrifugation at 20000 g for 90 min at 4 °C and suspended with 1× Laemmli buffer (diluted with RIPA buffer). After mixing with Laemmli buffer, the cell lysates and VLPs were incubated at 95 °C for 10 min. 12% or 15% acrylamide gels were used for SDS-PAGE. When CD81 was detected, 15% gel was used. The proteins were transferred to PVDF membranes, followed by blocking with 5% milk in TBS for at least 1 h. The membranes were incubated with primary antibodies at 4 °C o/n or at RT for 2-3 h and washed with TBS-T (0.1% Tween-20 in TBS) 3× 10 min. Secondary antibodies were incubated with the membranes in the dark for 1 h at RT and washed with TBS-T three times. Detection was performed using the Odyssey Fc Imaging System.

### trVLP luciferase assay

The cells were lysed with 200 µl luciferase lysis buffer (25 mM glycylglycine (pH 7.8), 15 mM MgSO_4_, 4 mM EGTA, 10% (v/v) glycerol, 1% (v/v) Triton X-100, 1 mM DTT (before use) in H_2_O) at RT for 15 min. For firefly luciferase (Fluc) assay, 40 µl Fluc assay buffer (0.1 M KH_2_PO_4_ /K_2_HPO_4_ pH 7.8, 15 mM MgSO_4_, 5 mM ATP in H_2_O) was added to opaque white 96-well plates and mixed with 40 µl cell lysates and 40 µl Fluc substrate buffer (0.28 mg/ml D-Luciferin in Fluc assay buffer). For NanoLuc luciferase (Nluc) assay, Nluc substrate buffer was prepared by diluting Nluc substrate in Nluc assay buffer at a ratio of 1/50 according to the manufacture’s protocol (Nano-Glo Luciferase Assay System). 40 µl cell lysate was added to opaque white 96-well plates and mixed with 40 µl Nluc substrate buffer. The measurements were done with the Cytation3 Cell Imaging Multi-Mode Reader.

### NFκB luciferase assay

A luciferase reporter plasmid was used for analysis of NFκB activity, followed by firefly luciferase signal detection using the luciferase assay system of Promega (#E1500). HEK293T cells seeded in a 96-well format (25000 cells/well) were transfected with 0.1 µg of NF-κB reporter plasmid (pGL3-NF-κB F-Luc), 0.05 µg of potential NFκB activator (EBOV proteins or constitutively active mutant of IKKβ) and, for the CD81 titration experiment, an additional of 0.1 µg of control or CD81-encoding pWPI plasmid. 24 h.p.t., cells were lysed with 40 µl of 1x Cell Lysis Buffer (diluted in H_2_O) for 5 min at RT. 30 µl of cell lysate was mixed with 30 µl luciferase substrate in a white 96-well plate and relative light units per second (RLU/s) were measured in the Berthold microplate reader.

### RT-qPCR

RNAs were extracted from the cells with RNeasy Mini Kit with on-column DNase digestion (Qiagen). 1 µg RNA was used for reverse transcription with QuantiTect Reverse Transcription kit (Qiagen). For transcription of EBOV minigenome vRNA and cRNA, specific primers were used, vRNA: ATTGAAGATTCAACAACCCTAAAG, cRNA: AATATGAGCCCAGACCTTTCG) (78). Oligo(dT) (Thermo Fisher) was used for reverse transcription of mRNA. After reverse transcription, the reaction mixture (20 µl) was diluted with 40 µl RNase free water, and 5 µl diluted reverse transcription mixture was used for qPCR. qPCR reactions were prepared with Luna Universal qPCR Master Mix (New England Biolabs) and 0.3 µM (final concentration) primers (Table 3). Measurements were done with LightCycler 480 System. qPCR results were analyzed using the ΔΔCp method (79).

**Table 3.**
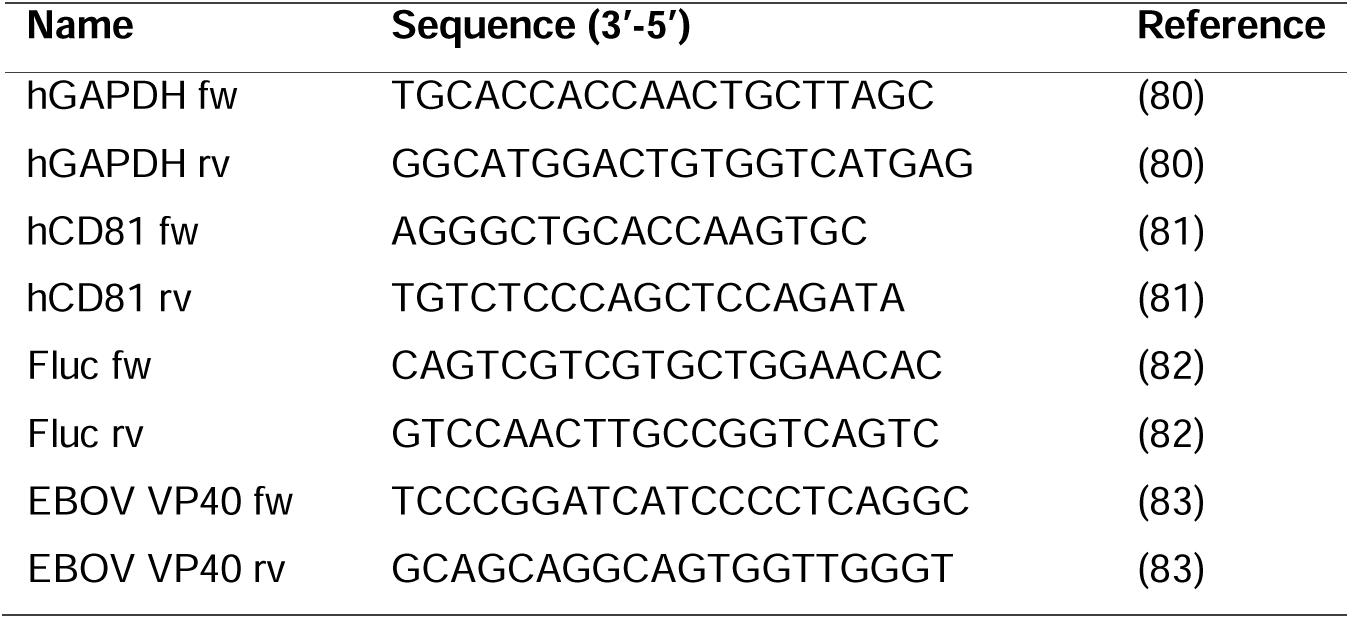
Primers for qPCR.

### BiFC assay

Kusabira-green (KG) based bimolecular fluorescence complementation (BiFC) assay from MBL life science (CoralHue® Fluo-chase Kit, a gift from AG Sauter) was used to analyze protein-protein interaction. HEK293T cells in 96-well plates were transfected with 75 ng KGN expression plasmids and 75 ng KGC expression plasmids. In the assay to analyze whether CD81 interacts with NP in the presence of VP35, 50 ng KGN expression plasmids and 50 ng KGC expression plasmids were transfected together with 50 ng pCAGGS-VP35_v1.1. 2 days post transfection, the cells were harvested and surface stained with PE-conjugated CD81 antibody for flow cytometry.

### Production and infection of lentivirus pseudotypes

To produce EBOV GP or VSV G pseudotyped GFP expressing lentiviruses, HEK293T cells in 6-well plates were transfected with 1.125 µg psPAX2, 1.5 µg pWPXLd and 0.125 µg pVSV-G or pCAGGS ZEBOV GP wt using PEI. After incubation o/n, the medium was changed and medium containing lentivirus pseudotypes was harvested 3 days later. The lentivirus-containing medium was cleared by centrifugation at 3200 g for 10 min. 0.9 ml of cleared medium was used for infection of HEK293T cells in 12-well plates and the medium was changed one day post infection. 3 days post infection, the cells were harvested and fixed with 2% PFA for flow cytometry.

### EBOV trVLP assay

The EBOV trVLP production and infection assay was performed according to (63). HEK293T cells in 12-well plates were transfected according to Table 4 scheme 1 to produce trVLP-nluc and trVLP-dGP-nluc using PEI, and the medium was changed after o/n incubation. To produce trVLP-nluc and trVLP-dGP-nluc in Huh7.5 cells, half the amount of the plasmids (Table 4 scheme 2) was transfected using jetPRIME, and the medium was changed 4 hours post transfection. 3 d.p.t., the cells were harvested for WB. In CD81 restoration experiments, HEK293T CD81 KO cells in 12-well plates were pre-transfected with pWPI-hCD81-HAHA-BLR (0, 0.0625, 0.125, 0.25, 0.5 and 1 µg) together with pWPI (1, 0.9375, 0.875, 0.75, 0.5 and 0 µg) in a total of 1 µg plasmids one day before transfection to produce trVLPs. Table 4 scheme 2 and 0.05 µg pCAGGS-luc2 (transfection ctrl) were transfected to produce trVLP-nluc or trVLP-dGP-nluc, half of the cells were harvested for WB and half of the cells were prepared for luciferase assay (3 d.p.t.). Amounts indicated in table 5 scheme 2 were transfected to produce trVLP-GFP or trVLP-dGP-GFP, and the cells were harvested for WB (3 d.p.t.). For live cell imaging experiments, cells in 96-well plates (triplicates) were transfected with 1/10 of scheme 2 (Table 5) together with 0.05 µg (10×) mCherry-C1 (transfection control). Live cell imaging started 6 h post transfection after medium change. For qPCR experiments, Table 4 scheme 1 and 0.1 µg pCAGGS-luc2 were transfected, and the cells were harvested 3 days post transfection.

**Table 4.**
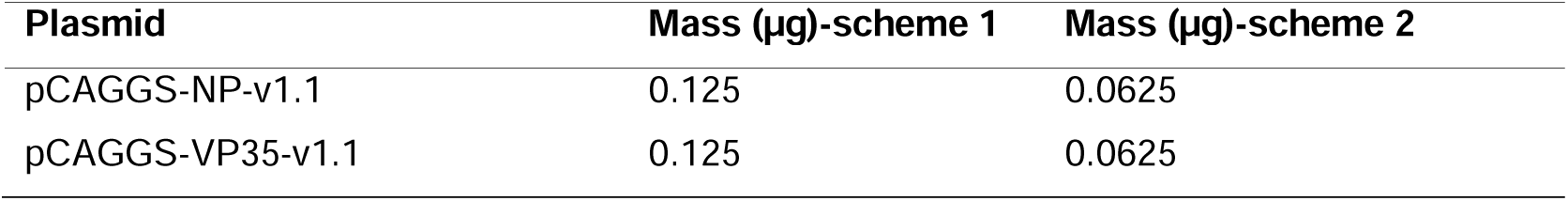
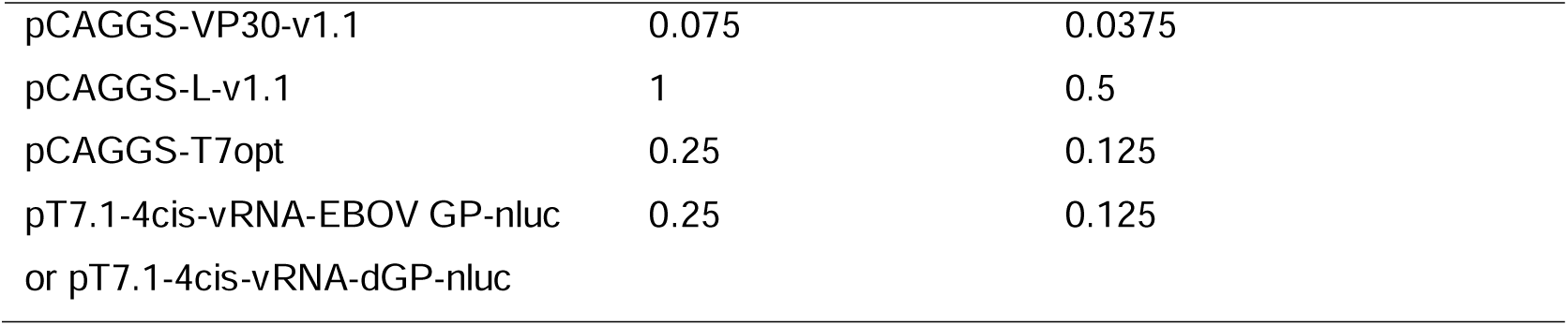
Transfection scheme to produce trVLP-nluc and trVLP-dGP-nluc.

**Table 5.**
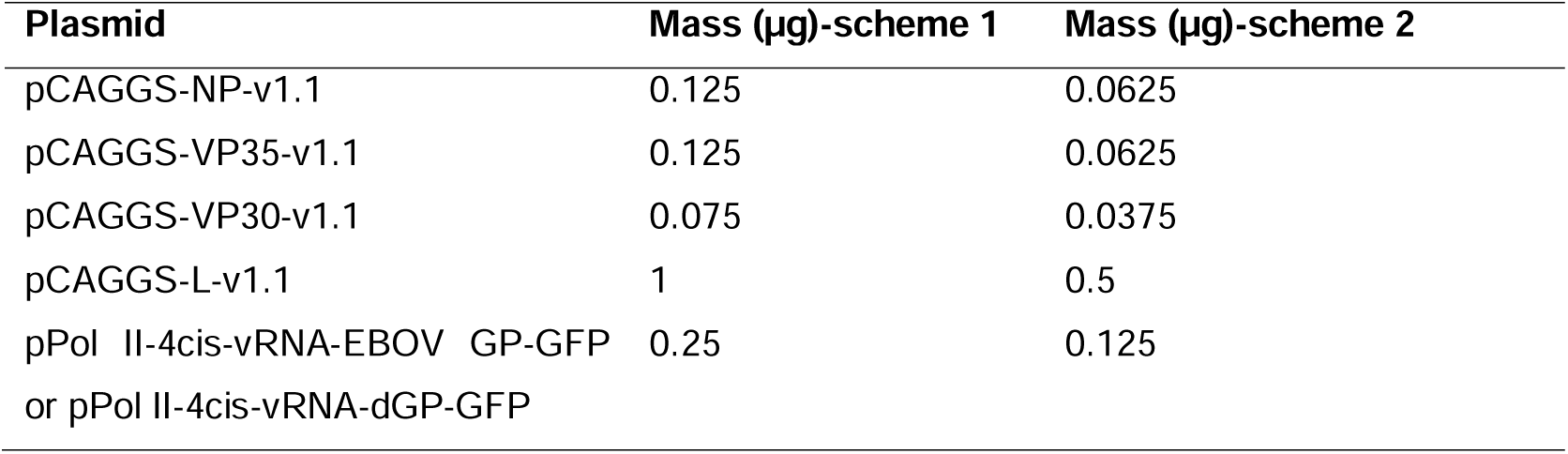
Transfection scheme to produce trVLP-GFP and trVLP-dGP-GFP.

To produce trVLP-GFP for infection, HEK293T cells in 6-well plates were transfected as indicated in Table 5 scheme 1 (pPol II-4cis-vRNA-EBOV GP-GFP). After incubation o/n, the medium was changed and trVLP-GFP containing medium was harvested 3 days later. The trVLP-containing medium was clarified by centrifugation at 800 g for 5 min and supernatants were used for infection. The produced trVLPs were passaged twice and used for infection of target cells in the experiments. To produce trVLP-dGP-GFP pseudotyped with EBOV GP or VSV G (trVLP-dGP-GFP_EBOV GP or VSV G) for infection, HEK293T cells in 6-well plates were transfected with 2× scheme 1 (pPol II-4cis-vRNA-dGP-GFP) and 0.125 µg (2×) pCAGGS ZEBOV GP wt or pVSV-G. After incubation for 4 h or o/n, the medium was changed and VLP containing medium was harvested 3 days later. The VLP-containing medium was centrifuged at 3200 g for 10 min and supernatants were taken for infection. For trVLP-GFP infection, HEK293T cells in 12- well plates were pre-transfected as scheme 1 (Table 6) one day before infection. 1 ml of cleared trVLP-GFP containing medium was added to the cells and medium was changed 1 day later. 3 days post infection, the cells and medium were harvested. 1/5 cells were collected for flow cytometry, and the remaining cells and medium were prepared for WB. For infection with trVLP-dGP-GFP_EBOV GP or VSV G (Fig. 5C), HEK293T cells in 12-well plates were pre-transfected with EBOV ribonucleoprotein complex (RNP) (NP, VP35, VP30 and L) and Tim1 as indicated in scheme 1 (Table 6) and 0.1 µg pCAGGS-luc2 (transfection ctrl, optionally) was co-transfected 1 day before infection. The cells were infected with 0.9 ml of cleared trVLP-dGP-GFP_EBOV GP or VSV G for 1 day and the medium was changed. 3 days post infection, the cells were harvested for flow cytometry. For infection of HEK293T cells with trVLP-dGP-GFP_EBOV GP with HEK293T wt cells (Fig. 5E), the cells in 96-well plates (triplicates) were pre-transfected with 1/10 scheme 2 (Table 5) + 0.05 µg (10×) pmtagBFP-C1 (transfection control) + 0.5 µg (10×) pWPI-hCD81-HAHA-BLR or pWPI_BLR. In case of Tim1-, pCAGGS-MCS instead of pCAGGS-Tim1-v1.2 was transfected. 1 day later, the cells were infected with 100 µl clarified trVLP-dGP-GFP_EBOV GP for 1 day, followed by medium change. 2 days post infection, the cells were harvested and surface stained with PE-conjugated CD81 antibody for flow cytometry. In the CD81 antibody treatment experiment, HEK293T cells seeded in 96-well plates (triplicates) were pre-transfected 1/10 of scheme 2 (Table 20) and 0.05 µg (10×) mCherry-C1. 1 day later, the medium was removed from the cells and 60 µl medium containing no antibody, 10 or 2 µg/ml anti-human CD81 Antibody (5A6) or Mouse IgG1, κ Isotype Ctrl Antibody was added. 1 h later, the cells were infected with 60 µl cleared trVLP-dGP-GFP, trVLP-dGP-GFP_EBOV GP or VSV G. 2 days post infection, the cells were harvested for flow cytometry. For flow cytometry, the cells were either fixed with 2% PFA or directly measured.

**Table 6.**
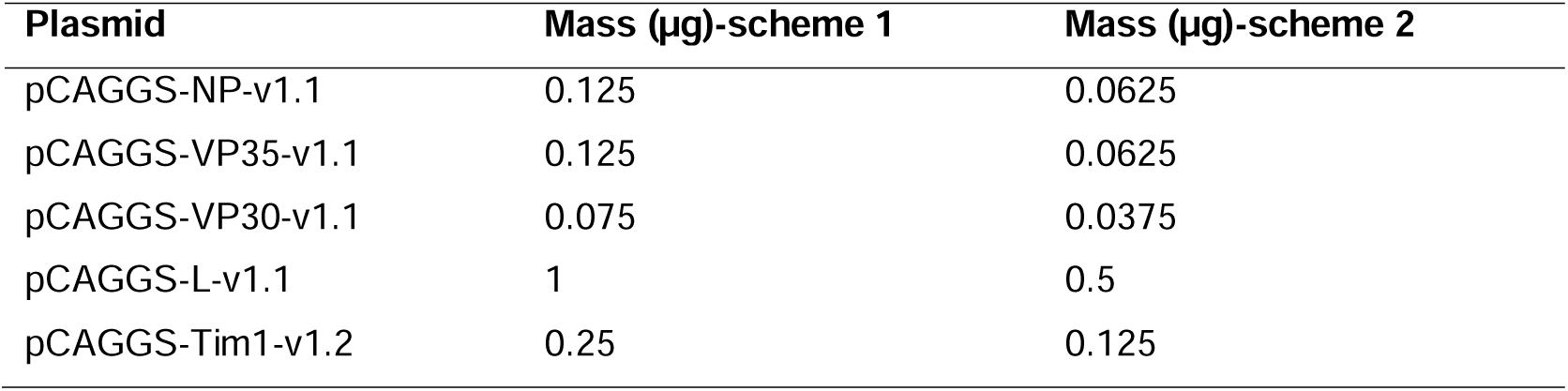
Transfection scheme for trVLP infection.

### EBOV infection

HEK293T control cells in 6-well plates were infected with rgEBOV-eGFP (84) at a MOI of 1. Monocyte-derived macrophages (MDM) were generated as described in cell culture section and seeded in 6 well plates with a density of 9×10^4^ cells per well. MDMs were infected the following day at a MOI of 0.5. 24 h post infection (h.p.i.) (HEK293T) or 6 d.p.i. (MDM), the cells were harvested using Accutase (BioLegend) and fixed with 4% PFA at 4 °C for at least 24 h or using BD Cytofix Fixation Buffer (BD Biosciences) at room-temperature for at least 1 h. Subsequently, the cells were washed with PBS and stained with PE-conjugated antibodies against CD81, CD63 and CD9 for flow cytometry. In the experiment with the CD81 antibody (5A6) treatment, HEK293T cells in 12-well plates were treated with no antibody (medium only), 5 or 1 µg/ml anti-human CD81 Antibody (5A6) or Mouse IgG1, κ Isotype Ctrl Antibody. 1 h later, the cells were infected with rgEBOV-GFP at a MOI of 0.3 or 1, with the virus being diluted in medium with or without antibody accordingly. 24 h post infection, the cells were harvested and fixed with 4% PFA at 4 °C for at least 48 h for flow cytometry. All HEK293T infection experiments with EBOV were performed in the BSL4 laboratory of the Friedrich-Loeffler-Institute following approved standard operating procedures. MDM infection experiments were performed in the BSL4 laboratory of the Bernhard-Nocht-Institute for tropical medicine following approved standard operating procedures. Antibodies used for the respective staining of EBOV infected MDM are detailed in Table 2 (FC MDM). Blood samples for MDM generation from 8 donors were obtained from a cohort of healthy volunteers at the Bernhard Nocht Institute for Tropical Medicine (BNITM) in Hamburg (Germany) with approved consent. The protocol for the use of these samples was authorized by the ethics committee of the German Medical Association and the ethics committee of the Ärztekammer Hamburg (PV4780)

### Measles virus (MeV) infection

HEK293T control, CD81 KO and HeLa cells were seeded in a 12-well plate format at a density of 1.5×10^5^ (HEK293T) or 3×10^5^ (HeLa) cells per well. The following day they were infected with MeV-GFP (kindly provided by Ulrich Lauer, Virotherapy Center, Tübingen) diluted in Opti-MEM at a MOI of 0.1 or 1. A medium change was performed 3 h.p.i. and the cells were incubated for another 48 hours at 37°C. The infected cells were harvested, fixed with 2% PFA for 15 min at 37°C, washed with PBS and measured using the flow cytometer. For tetraspanin modulation, cells were stained with PE-conjugated antibodies against CD81, CD63 and CD9. In order to assess the effect of CD81 (5A6) antibody treatment, cells were treated one hour prior to infection with 1 µg/ml anti-human CD81 Antibody (5A6) or Mouse IgG1, κ Isotype Ctrl Antibody or left untreated.

### Software and analysis tools

FlowLogic (Inivai) and FlowJoc.10 was used for flow cytometry data analysis. Image Studio (LI-COR) was used for WB signals quantification. Incucyte (Sartorius) was used for quantification of microcopy images. ImageJ was used for IB and p65 quantification of immunofluorescence microscopy images. Microsoft Excel and Graphpad Prism 9.4.1 was used for statistical analysis and Corel Draw for generation of figures.

## Results

### EBOV GP reduces cell surface expression levels of tetraspanins CD81, CD63 and CD9

To identify EBOV GP-modulated cell surface receptors, a flow cytometry-based screen was performed. HeLa cells were transfected to express EBOV GP together with GFP via a bicistronic mRNA allowing to specifically identify cells that express GP (GFP-positive cells) and compare them to non-GP (GFP-negative) expressing cells in the same measurement (Fig. S1). Importantly, high GP-levels might induce cytotoxicity and cell death that could compromise our results. However, we did not detect altered forward or sideward scatter in our flow-cytometry plots upon GP-expression or increased AnnexinV-staining, as a proxy for apoptosis (Figure S1). Next, cells were stained with PE-labeled antibodies against 332 plasma membrane proteins in a 96-well format and subjected to medium-throughput flow cytometry-measurement. GP-mediated fold of surface protein downmodulation was calculated by dividing the MFI (mean fluorescence intensity) of the non-GP (GFP-negative) population by the MFI of the GP-expressing (GFP-positive) cells. EBOV GP modulated surface proteins were identified according to a threshold of modulation >2-fold (Table S1) and 21 receptors (Figure 1a) were selected for further validation in HeLa and HEK293T cells (Figure 1b, Table S2). HLA, CD81, CD63, CD9, CD201 (endothelial protein C receptor), CD184 (C-X-C chemokine receptor type 4) and CD59, showed reduced cell surface levels upon EBOV GP-expression in both cell lines (Figure 1b and Figure S2a-b). HLA was reported before to have lower accessibility to antibodies due to shielding by EBOV GP glycans (60). Hence, reduced cell surface detection could be due to shielding of receptors or their removal and internalization, i.e. active downregulation. We next decided to focus on CD81, CD63 and CD9, since these three proteins belong to the tetraspanin family and were all affected by GP (Figure 1a-c). Reduced cell surface levels of these three tetraspanins was confirmed in the context of authentic EBOV infection in which both surface and total cellular CD81, CD63 and CD9 were affected (Figure 1d). Strikingly, lower levels of CD81 and to a lesser extent CD9 were also detected at the cell surface of primary human monocyte-derived macrophages (MDM) infected with authentic EBOV (Figure 1d and f). In order to distinguish general viral effects from EBOV-specific mechanisms, measles virus (MeV) infection was included as a control throughout this study. Like EBOV, MeV is an enveloped, negative-sense RNA virus, but it employs distinct entry and release mechanisms (85). The tetraspanins were not modulated upon Measles-Virus (MeV)-infection (Figure 1e and f). Taken together, EBOV GP modulates the tetraspanins CD81, CD63 and CD9 in various cell lines as well as in the context of authentic EBOV infection in primary macrophages.

### Tetraspanin modulation is specific for filoviral GPs

To determine whether the tetraspanins are also affected by other filoviruses, 293T cells were transfected to express GP of Marburg virus (MARV), Sudan virus (SUDV), Reston virus (RESTV) and Taï Forest virus (TAFV). All filoviral GPs were able to reduce CD81, CD63 and CD9, suggesting that tetraspanin modulation is a conserved feature of various members of the *Filoviridae* family (Figure 2a). Next, we characterized GP1/2 domains (Figure 2b) and several GP mutants (Figure S2c). Neither GP1 nor GP2 alone induced significant tetraspanin modulation (Figure 2b). Notably, GP2 was not detectable by Western blot when expressed individually, and therefore no conclusion can be drawn here regarding its potential to modulate tetraspanins (Fig. S2d). The GP mutants tested had altered residues in the receptor-binding site (RBD, located in GP1), the fusion loop, (FL, located in GP2) and the transmembrane domain (TM, located in GP2), all of which have been previously shown to be important for GP functionality (Figure S2c). All GP mutants still reduced CD81, CD63 and CD9 levels, although two mutations in the RBD (L111A, L122A), two mutants with altered residues in the FL (531A532A533A and 535A536A537A) and the ZLZ mutant (TM is changed to the TM of LASV GP) were less effective (Figure S2e). We further analyzed if non-filoviral GPs might dysregulate CD81, CD63 or CD9. For this, a set of previously characterized viral GPs were transfected to be expressed in 293T cells together with GFP and assessed for cell surface tetraspanin levels (Fig. 2c). Of note, apart from EBOV GP, none of the tested GPs affected any of the three tetraspanins, suggesting that modulation of CD81, CD63, and CD9 is a specific feature of filoviral GPs.

**Fig. 2.**
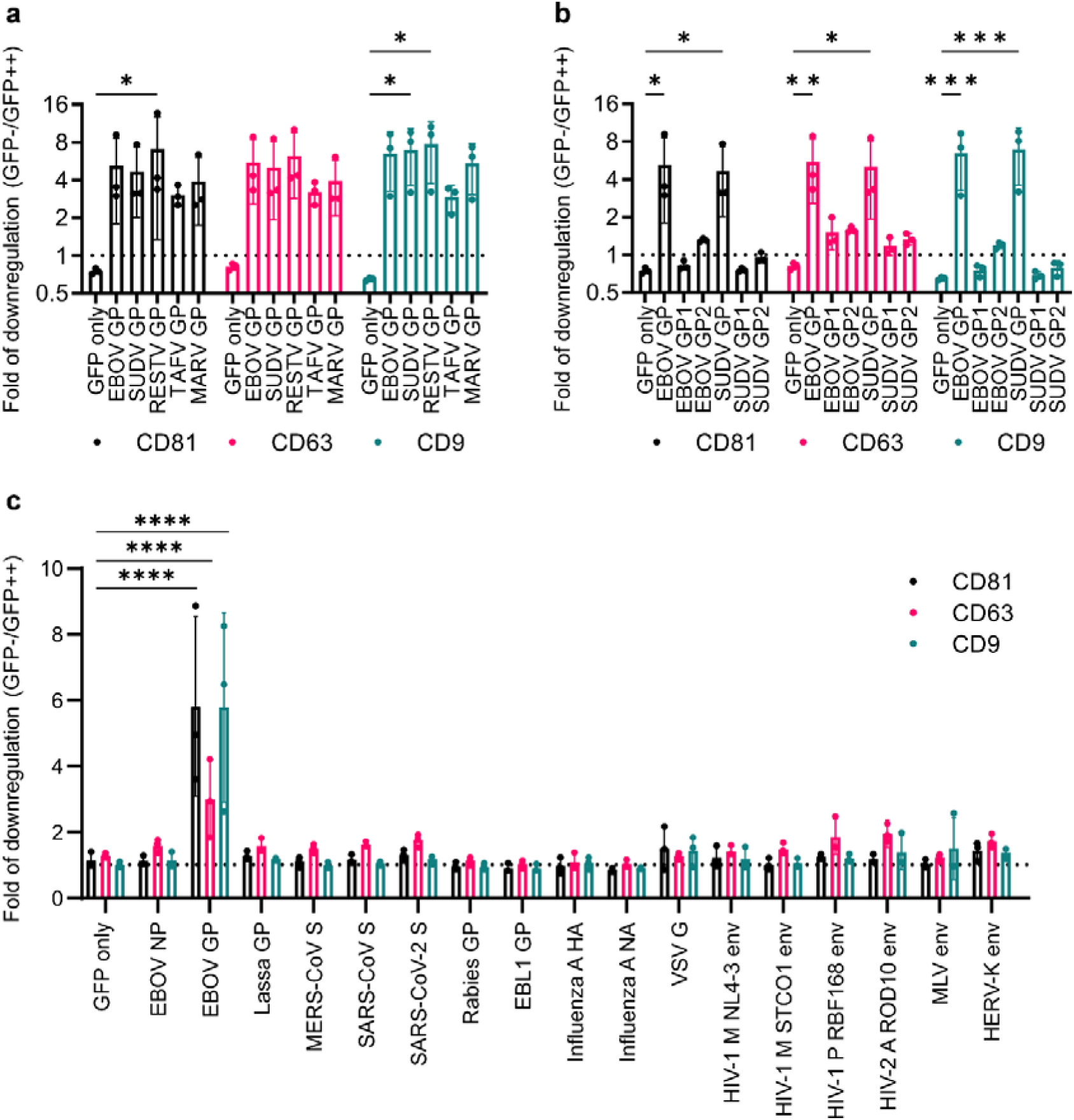
CD81, CD63 and CD9 downregulation is conserved among EBOV strains but not other viral glycoproteins. **(a-c)** 293T cells were transfected to express GFP only or the (a) indicated filoviral GPs, (b) GP subunits or (c) viral glycoproteins together with GFP. 2 d.p.t., the cells were harvested and analyzed for surface expression of CD81, CD63 and CD9 by flow cytometry. Indicated are mean values of fold of modulation (GFP-/GFP++) +/- SD (n=3). Two-way ANOVA with Dunnett correction was used for statistical analysis: <0.05 (*), <0.01 (**), <0.001 (***), <0.0001 (****).

### CD81, but not CD63 and CD9, suppresses EBOV trVLP replication

To reveal why EBOV interferes with CD81, CD63 and CD9, the EBOV trVLP assay was used (63), as this system allows to analyze and model the different aspects of the EBOV life cycle. 293T and Huh7.5 control, CD81 KO, CD63 KO and CD9 KO cells were generated using CRISPR/Cas9 and KO confirmed by flow cytometry (Figure S3a and b). 293T control and KO cells were transfected to express EBOV RNP (NP, L, VP35 and VP30), a tetracistronic (4cis) minigenome encoding a nanoluciferase (nluc) reporter as well as the other three EBOV proteins (GP, VP40 and VP24) and a plasmid expressing T7 RNA polymerase, mediating the initial transcription of the minigenome. Expression levels of the EBOV matrix protein VP40, encoded by the minigenome and therefore reflecting EBOV minigenome replication and transcription, were analyzed in both cell lysates and culture supernatants (medium) by Western blot. WT and a GP-deleted (dGP) minigenome were used to compare the effect of CD81, CD63 and CD9 on trVLP replication with and without GP expression.

KO of CD81, but not CD63 and CD9, led to significantly higher VP40 levels in cells and supernatants of trVLP-dGP producing cells. Similar results were observed in cells producing WT trVLP, suggesting a negative role of CD81 in EBOV trVLP viral RNA synthesis and/or protein expression (Figure 3a). Besides, KO of CD81 resulted in higher increase in VP40 levels in cells (and supernatants) producing trVLP-dGP (∼4,19) as compared to wildtype trVLP (∼2,87), indicating that expression of EBOV GP can counteract the inhibitory effect imposed by CD81 (Figure 3a). Furthermore, VP40 levels in cell culture supernatants were normalized to VP40 in cell lysates, indicating that KO of CD81 (as well as CD63 and CD9) has no measurable effect on trVLP release in this system (Figure 3a). The aforementioned results were recapitulated and confirmed independently in Huh7.5 hepatocellular carcinoma cells (Fig. 3b) and with a employing a GFP-encoding trVLP construct, monitoring virus production over several days using GFP as a proxy (Figure S3c). In both cell lines, albeit more pronounced in Huh7.5 cells, KO of CD81 increased GFP expression over time especially in the absence of GP (trVLP-dGP-GFP), likely due to the absence of CD81 antagonism (Figure S3c). In general, VP40 and reporter gene expression was higher in trVLP-dGP producer cells than trVLP-WT producer cells, which might be related to the inverse correlation between minigenome length and efficiency of viral protein expression (Fig. S3d and (63)). Altogether, the data indicates that CD81, but not CD63 and CD9, suppresses EBOV trVLP replication in 293T and Huh7.5 cells.

**Fig. 3.**
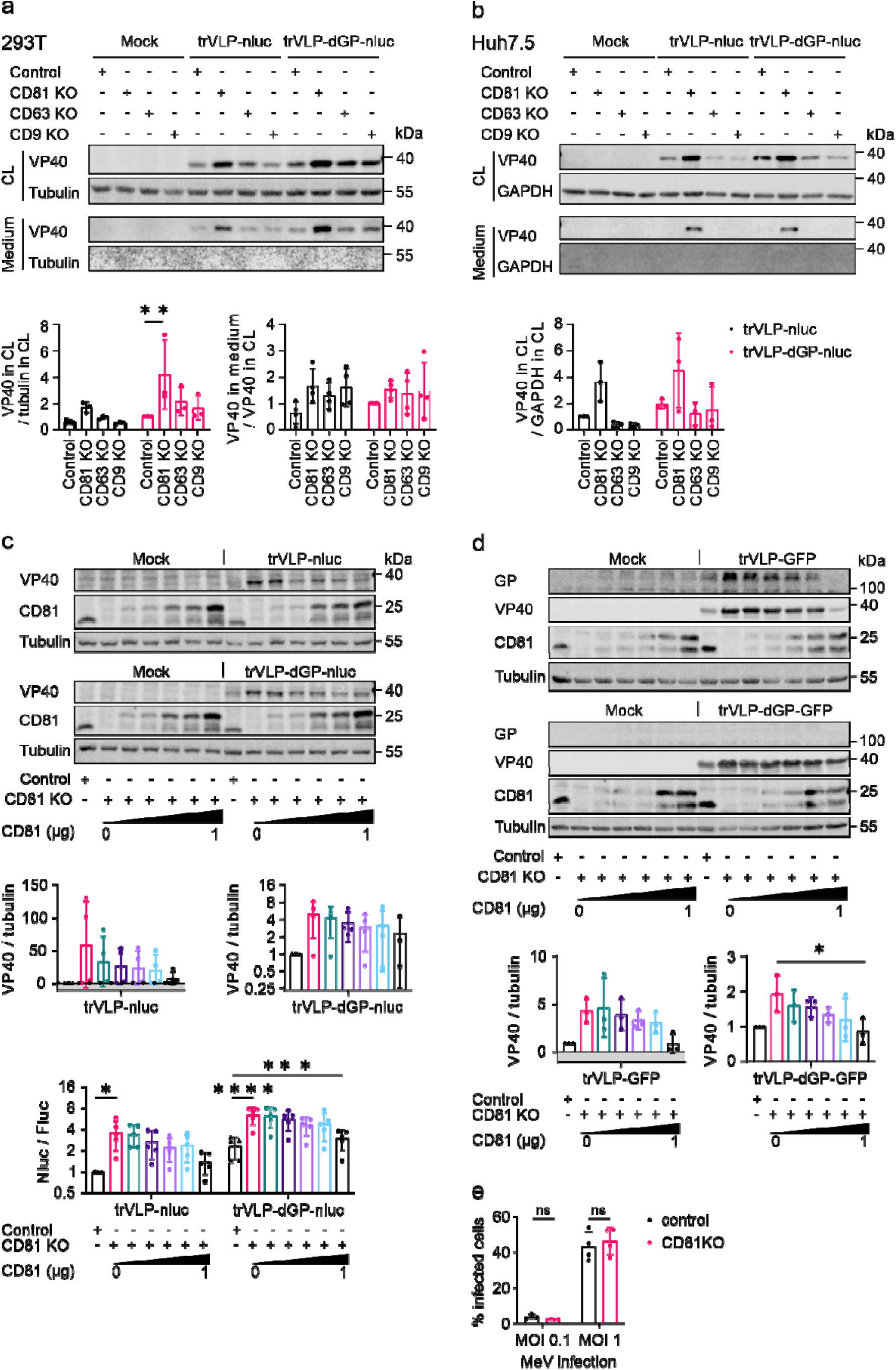
CD81, but not CD63 and CD9, suppresses EBOV trVLP replication. In order to assess EBOV replication and transcription, cells were transfected with RNP expression plasmids and a tetracistronic minigenome plasmid encoding nLuc or GFP and EBOV VP40, GP and VP24 (trVLP) or a GP deleted minigenome plasmid (trVLP-dGP). (a) 293T and (b) Huh7.5 control, CD81 KO, CD63 KO and CD9 KO cells were transfected for trVLP-nluc/-dGP-nluc replication and production. 3 d.p.t., cells and medium were harvested for WB analysis (n=3). Indicated are representative WB images and mean values of WB signal quantification (normalized to control cells producing trVLPs, +/- SD). (c) 293T control and CD81 KO cells were mock or pre- transfected to express gradient amounts of CD81-HA. 1 day later, the cells were transfected to produce trVLP-nluc/-dGP-nluc. A Fluc expression plasmid was co-transfected as a transfection control for normalization. 3 d.p.t., an aliquot of the cells was harvested to analyze Nluc and Fluc activity, and the remaining cells were harvested for WB analysis (n=4-5). Indicated are mean values of WB signal quantification (normalized to control cells, +/- SD) and mean values of Nluc activity (normalized to Fluc and control cells producing trVLP-nluc, +/- SD). (d) 293T cells mock or pre- transfected to express gradient amounts of CD81-HA were transfected for trVLP-GFP and trVLP-dGP-GFP replication. 3 d.p.t., the cells were harvested for WB analysis. Indicated are mean values of WB signal quantification +/- SD (n=3). (e) As specificity control, 293T control and CD81 KO cells were infected with MeV-GFP. 2 d.p.i., infected (GFP+) cells were quantified by flow cytometry (n=4). Two-way ANOVA with Dunnett correction was used for statistical analysis: <0.05 (*), <0.01 (**), <0.001 (***), <0.0001 (****).

To confirm the suppressive role of CD81 in EBOV trVLP replication in another independent experimental setting, CD81 expression in 293T CD81 KO cells was restored by exogenously expressing increasing amounts of HA-tagged CD81 (CD81-HA). trVLP VP40 as well as nluc expression (reflecting viral RNA synthesis/gene expression) were analyzed by Western blot and luciferase activity assay. In accordance with our previous results, not only VP40 but also nluc expression increased upon KO of CD81 (Figure 3c). Furthermore, expression of CD81 reduced VP40 and nluc expression in a dose dependent manner in 293T CD81 KO cells (Figure 3c). To exclude the possibility that CD81 interferes with T7 polymerase mediated initial transcription of the 4cis-minigenome, a similar experiment was performed with an RNA polymerase II (pol II) driven 4cis-minigenome plasmid, which has similar gene organization as the T7 driven 4cis-minigenome. Similarly, KO of CD81 resulted in increased GP and VP40 expression, and exogenous expression of CD81 in CD81 KO cells reduced GP and VP40 expression (Figure 3d). Finally, we again included MeV as negative control, which showed unlike EBOV similar infection levels in CD81 KO, as compared to control cells (Figure 3e).

Altogether, the cumulated data assessing replication of various trVLP minigenomes in different cell systems with CD81 overexpression as well as KO conditions support a restricting role of CD81 in the EBOV life cycle, that is counteracted by GP.

### Mechanism of EBOV GP and VP40 mediated interference with CD81

Given that potent cellular restrictions to virus replication might be counteracted by more than one mechanism, we sought to determine whether EBOV employs, in addition to GP, other viral proteins to modulate CD81. 293T cells were transfected to express GFP only or co-express GFP and EBOV proteins. Cell surface as well as intracellular CD81 expression were analyzed by flow cytometry. Of note, CD81 was also downregulated by VP40, albeit less potently than GP, at both surface and intracellular levels in 293T cells (Figure 4a and Figure S4a). Since only CD81, but not CD63 and CD9, restricts EBOV trVLP replication, the potential role of CD81 in the downregulation of CD63 and CD9 by EBOV GP was investigated. CD63 and CD9 were reduced by EBOV GP regardless of CD81 expression, in both 293T and Huh7.5 cells (Figure S4b and c). Next, we aimed to assess if the viral proteins induce internalization and possibly degradation of cell surface CD81, or if CD81 might be shielded from antibody binding via the GP glycans. For this, 293T cells expressing EBOV GP or VP40 were treated with the proteasome inhibitor MG132 or the lysosome degradation inhibitor bafilomycin A1 (BafA1). Cell surface as well as total CD81 expression were analyzed by flow cytometry. MG132 and BafA1 slightly blocked GP mediated reduction of surface CD81 whereas VP40-mediated downregulation of surface CD81 was strongly blocked by MG132 and partially by BafA1 (Figure 4b, upper panel, see primary flow cytometry plots). Furthermore, MG132 and BafA1 partially restored total CD81 levels upon GP expression and had a marked inhibitory effect on the ability of VP40 to reduce total CD81 (Figure 4b. lower panel).

**Fig. 4.**
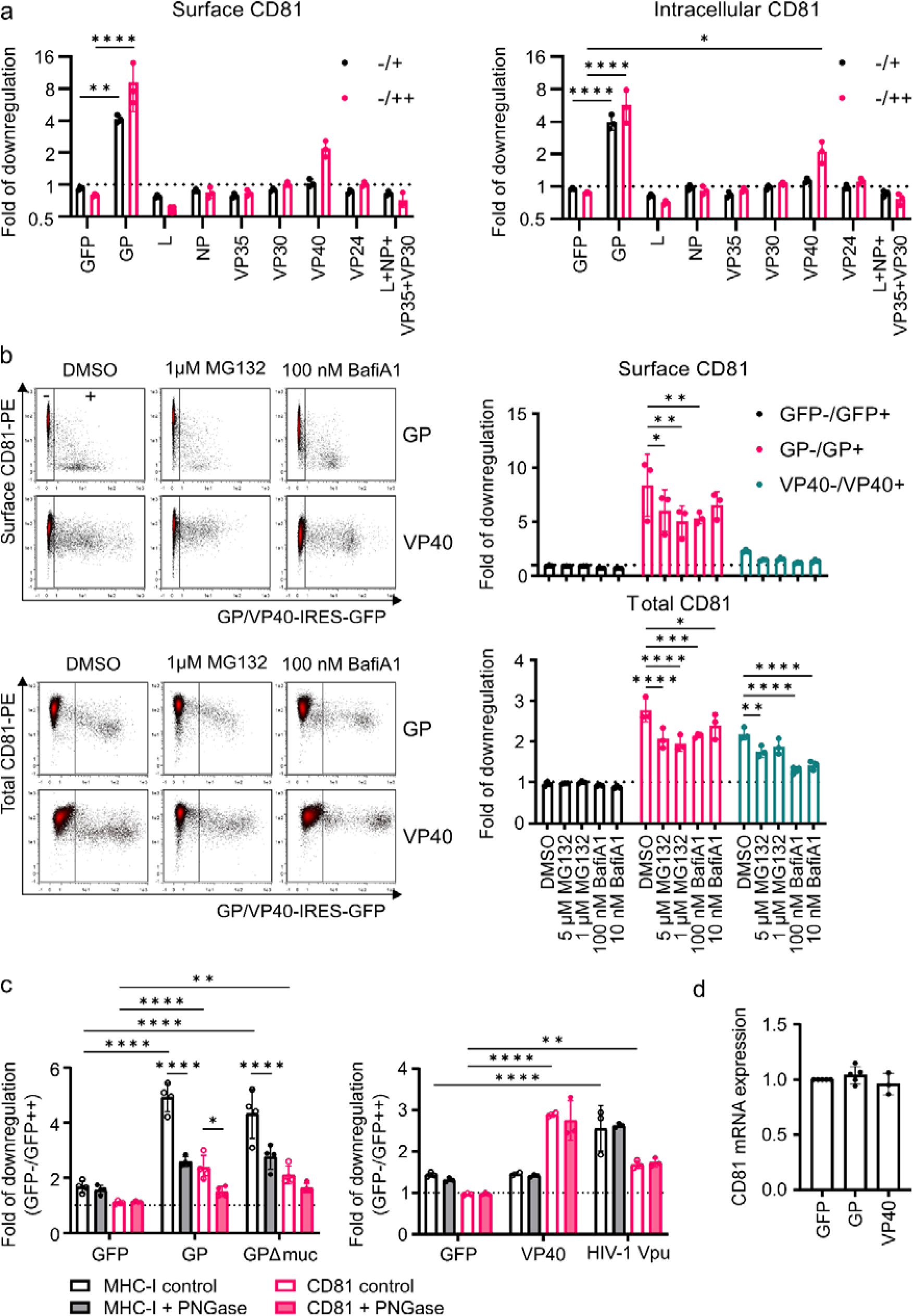
Mechanism of EBOV GP and VP40 mediated downregulation of CD81. (a) 293T cells were transfected to express GFP only or co-express GFP and EBOV proteins. 2 d.p.t., the cells were harvested for surface and intracellular CD81 staining and analyzed by flow cytometry. Indicated are mean values of fold of modulation +/- SD (n=3). Plotted are fold modulations by dividing MFIs of cells expressing medium (+) to high (++) levels of viral proteins with MFIs of cells that do not express the viral proteins (-) according to GFP-expression (see also Fig. S1) (b) 293T cells were transfected to express GFP only or co-expressing GFP and GP or VP40. 6 hours later, the cells were treated with DMSO only, MG132 or Bafilomycin A1. After 16 hours incubation, the cells were harvested for surface and total CD81 staining and analyzed by flow cytometry. Indicated are mean values of fold of modulation +/- SD (n=3) and representative density plots. (c) 293T cells were transfected to express GFP only, or GFP co-transfected with GP, a GP mutant that is lacking the mucin-like domain (GPΔmuc), VP40 or HIV-1 Vpu. 24 h.p.t. cells were left untreated or treated with PNGase (100 U/ml). 4 hours later the cells were harvested and analyzed for CD81 surface expression. Shown are mean values of fold of modulation +/- SD (n=4 left panel and n=3 right panel). (d) CD81 mRNA levels of GP or VP40-expressing 293T cells were quantified by RT-qPCR. Data was normalized to GAPDH and presented normalized to GFP-expressing cells (VP40 n=3) and (GFP/GP n=5). (a-d) Two-way ANOVA with Dunnett correction was used for statistical analysis: <0.05 (*), <0.01 (**), <0.001 (***), <0.0001 (****).

To assess the importance of the GP glycan shield for this phenomenon, we employed the GPΔmuc mutant that is devoid of the mucin-like domain, which is heavily glycosylated (86). On top, we treated GP-expressing cells with PNGase F to remove N-glycans before staining with CD81 antibodies or HLA-A (MHCI) as control (87, 88) (Fig. 4c). GPΔmuc reduced cell surface MHCI and CD81 slightly less efficient than WT GP. In addition, N-glycan removal by PNGase F partially restored cell surface antibody binding to MHCI and CD81 in GP-expressing cells (Fig. 4c, left panel). Of note, and as expected, when using non-glycosylated VP40 and HIV-1 Vpu, another tetraspanin-modulating viral protein (51), PNGase F treatment had no impact on modulation of MHCI or CD81 (Fig. 4c, right panel). Furthermore, neither GP nor VP40 transcriptionally impaired CD81 mRNA levels (Fig. 4d). In conclusion, CD81 surface and total expression levels are reduced via the concerted action of EBOV GP and VP40 involving pathways of proteasomal and lysosomal degradation, as well as CD81 surface shielding by GP.

### EBOV GP interacts with CD81

We next analyzed whether GP or VP40 interact with CD81 first using the Kusabira-green (KG) based BiFC assay (see M&M for details). CD81, GP and VP40 were expressed as fusion proteins with the N- or C- part of KG (KGN or KGC), and reconstitution of KG fluorescence can be measured as a proxy of protein interaction. Upon transfection of 293T cells to express the KGN/KGC fusions of GP and CD81, a strong increase in the percentage of KG expressing cells (KG+) could be measured, suggesting that both proteins interact (Figure 5a). However, whether CD81 interacts with VP40 remains unclear, due to the high background signal observed from controls (data not shown). To corroborate this result, we employed flow-cytometry based FRET that indicates close proximity of potential interaction partners at a distance below 10 nm (73, 89). We further expanded the analysis to GPs from other EBOV species and assessed FRET with the GP1 and GP2 subunits of EBOV (74). Importantly, GP2 fused to YFP is stably expressed (74). In line with the KG assay, all tested GPs exerted FRET with CD81, albeit with differential efficacy, and GP2 alone was sufficient to induce FRET (Fig 5b). Furthermore, proximity-ligation assay (PLA) supported close proximity of GP to endogenous CD81 in HeLa cells (Fig. 5c) which is corrobortated by areas of pronounced colocalization between CD81 and GP detected by immunofluorescence staining (Fig. 5d). Altogether, using various complementary methodology, the data supports a model in which GP directly interacts with CD81.

**Fig. 5.**
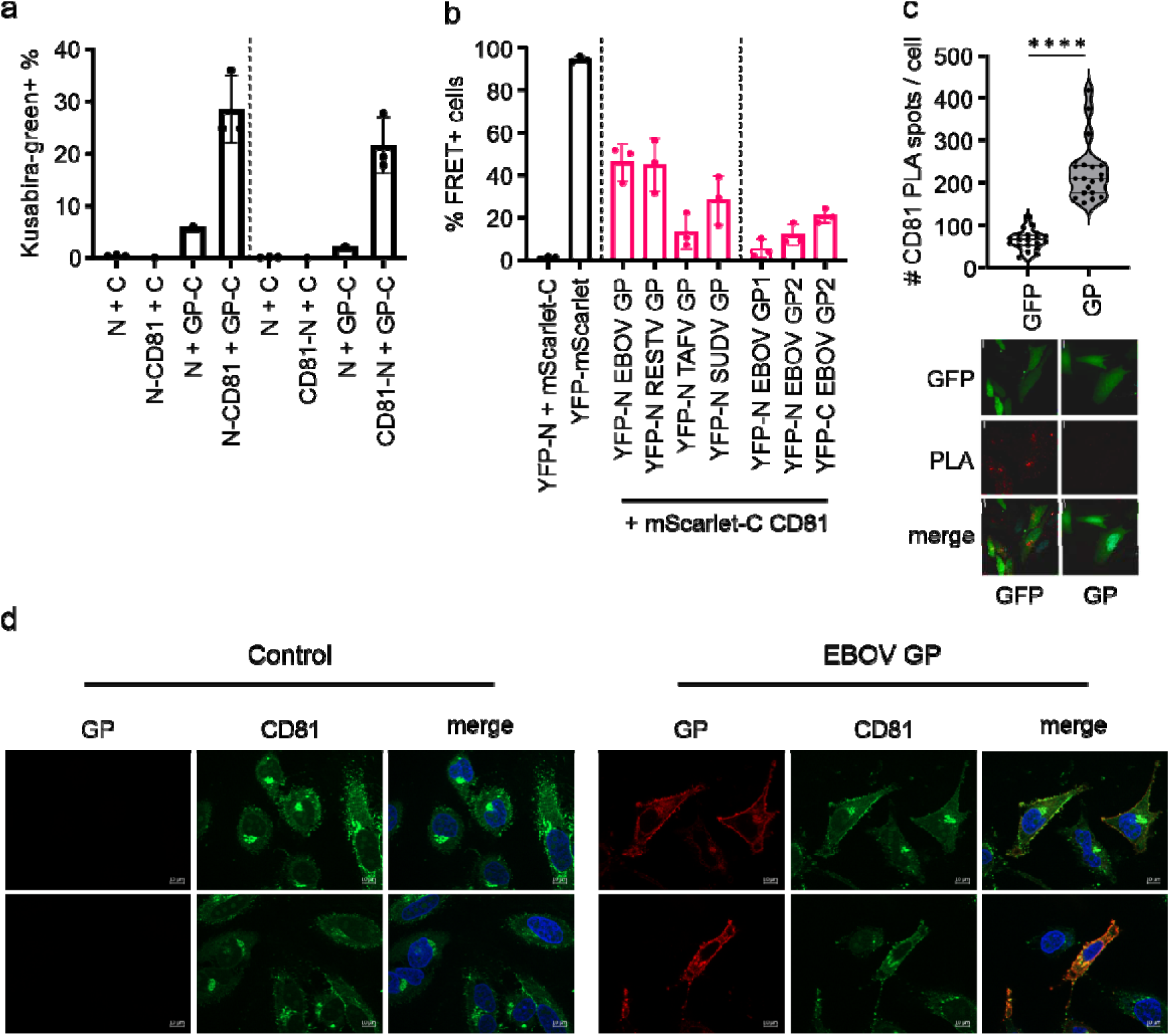
EBOV GP interacts with CD81. (a) 293T cells were transfected to express the N- and C- part of Kusabira-green as fusion protein with GP or CD81 as indicated and analyzed by flow cytometry. Mean values +/- SD (n=3). (b) Filoviral GPs fused to YFP were transfected together with a mScarlet-CD81 fusion protein in 293T cells. 24 h.p.t., cells were harvested and FRET signal was measured by flow cytometry. Indicated are mean values +/- SD (n=3) (c) HeLa cells were transfected to express GFP-only or GP together with GFP from a bicistronic mRNA. 24 h.p.t. cells were stained for PLA with CD81-specific antibodies as detailed in the M&M section. PLA signal from 20 cells from two independent transfections was analyzed for statistical assessment. Two-tailed non-parametric Mann-Whitney test: <0.0001 (****). (d) HeLa cells, mock-transfected or transfected to express GP were fixed 24 h.p.t. and stained for GP and CD81. Shown are representative images of 3 independent experiments.

### CD81 reduces overall levels of EBOV RNA-species and suppresses VP40-induced NF**κ**B activation

To elaborate which steps of EBOV trVLP replication are suppressed by CD81, we determined the effect of CD81 on EBOV trVLP mRNA, vRNA and cRNA synthesis in trVLP producer cells (90–92). Therefore, we measured RNA species in trVLP producer cells and found that KO of CD81 increased VP40 cRNA, and that there was also a trend to higher mRNA and vRNA levels (Figure 6a). cRNA and vRNA levels are dependent on viral genome replication, whereas mRNA levels are dependent on both mRNA transcription and the amount of vRNAs available as template for this process, and thus on both replication and transcription. This indicates that CD81 interferes with minigenome replication (as seen by the reduction of cRNA levels), whereas it remains unclear whether there is also a CD81-mediated suppression of transcription.

**Fig. 6.**
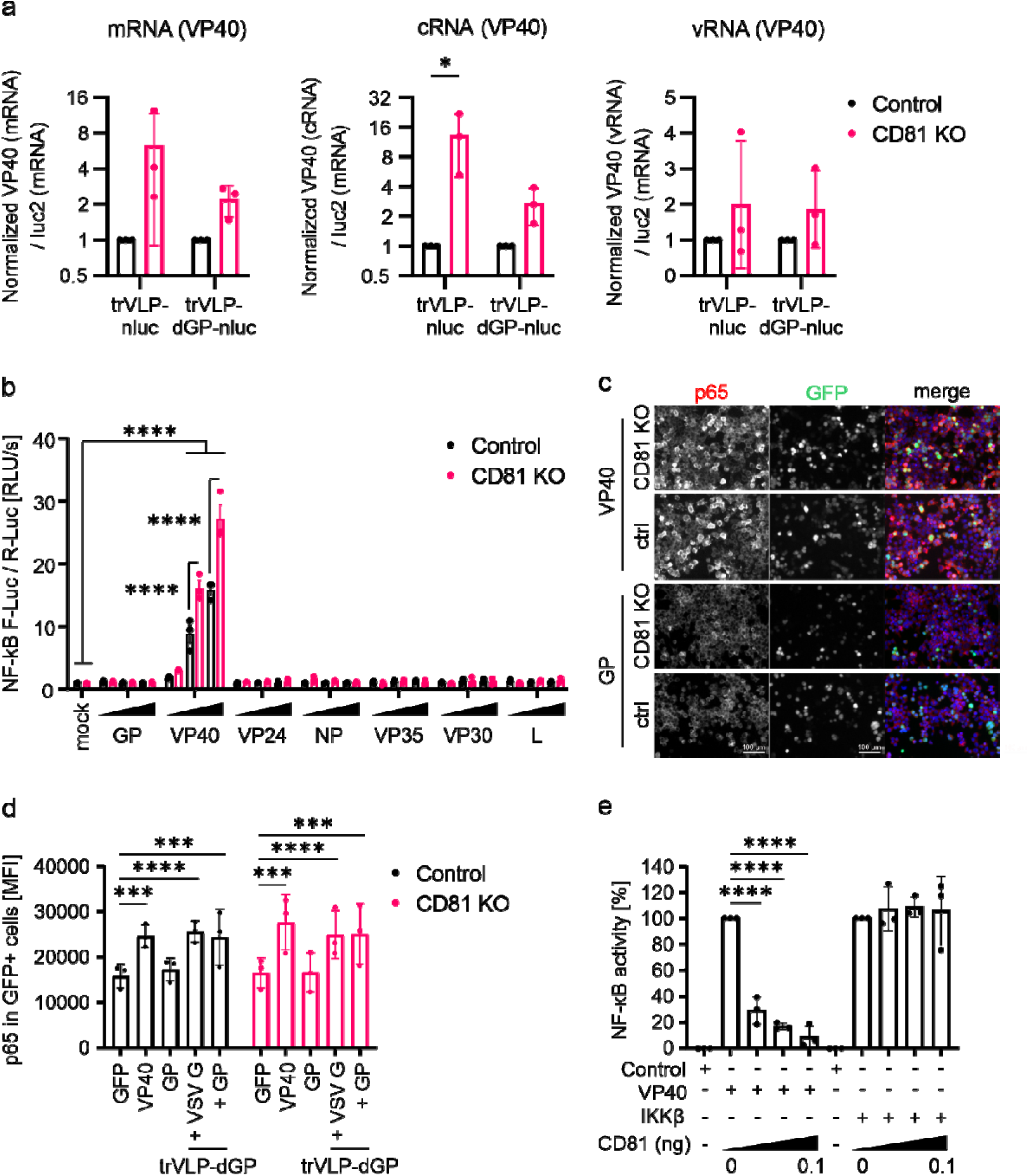
CD81 reduces levels of EBOV RNA-species and suppresses VP40-induced NFκB activation. (a) 293T control and CD81 KO cells were transfected for trVLP replication and transcription. 3 d.p.t., the cells were harvested for RNA extraction. vRNA, cRNA and mRNA levels of VP40 and the luc2 (fluc) mRNA level were determined by RT-qPCR. Indicated are mean values (normalized to luc2 and further the control cells) +/- SD (n=3). (b) 293T control and CD81 KO cells were transfected to express increasing amounts of EBOV proteins, a reporter expressing Firefly-luciferase (F-Luc) under control of a NFκB responsive promoter and Renilla-luciferase for normalization (R-Luc). 24 h.p.t cells were harvested, lysed and subjected to luciferase measurement. Indicated are mean F-Luc values (normalized to R-Luc) +/- SD (n=3). (c) Similar setup as in (b), however 293T cells were analyzed by fluorescence microscopy and endogenous p65 was stained by IF, nuclei with DAPI and EBOV protein expressing plasmids co-express GFP via an IRES. (d) Quantification of p65 mean fluorescence intensity in GFP-positive 293T cells expressing the indicated EBOV proteins (c) or previously infected with VSV G- or GP- pseudotyped EBOV trVLPs (sup. Figure S5). Indicated are mean values,+/- SD (n=3). (e) 293T cells were transfected to express VP40 or a constitutively active mutant of IKKβ and a F-Luc NFκB reporter. Furthermore, a control vector or increasing concentrations of a CD81-expressing vector were transfected. Luciferase reporter activity was measured 24 h.p.t. Indicated are mean values, background subtracted and normalized to corresponding 0 ng CD81 condition, +/-SD (n=3). (a) Two-way ANOVA with Šidák multiple comparison was used for statistical analysis: <0.05 (*), <0.01 (**), <0.001 (***), <0.0001 (****). (b,d,e) Two-way ANOVA with Dunnett’s multiple comparison was used for statistical analysis: <0.001 (***), <0.0001 (****).

EBOV genome replication and transcription take place in EBOV IBs; a process requiring specific viral protein interactions (31, 33). In this aspect, we sought to determine whether CD81 modulates EBOV IB formation. As previously described, sole expression of NP allows the formation of dot-like structure of IBs, and VP35 interacting with NP contributes to IB formation and viral RNA synthesis (93, 94). Additionally, VP35 is a chaperone of NP regulating NP homo-oligomerization and NP-RNA binding (95, 96). To investigate whether CD81 affects NP homo-oligomerization and NP-VP35 interaction, we again used the Kusabira-green based BiFC assay. EBOV NP and VP35 were expressed as fusion proteins with either KGN or KGC, and reconstruction of KG represents protein oligomerization or interaction. KO of CD81 did not result in increased number of cells showing NP, VP35 or NP to VP35-interaction (KG+ %)(Figure S5a). This indicates that CD81 does not have an impact on IB formation, which we independently confirmed by quantifying IB-formation in cells expressing the RNP proteins, or in RNP-expressing cells infected with trVLPs (Figure S5b and c).

We recently demonstrated that CD81 can suppress NFκB activation (46) and therefore hypothesized that NFκB is dysregulated by viral proteins in trVLP infected cells, possibly in a CD81-dependent manner. To test for this, we first transfected 293T cells to express various EBOV proteins and analyzed NFκB-dependent luciferase reporter activity. Of note, VP40 expression resulted in a strongly increased luciferase signal as a proxy for NFκB activation and this effect was even more pronounced in CD81 KO cells (Figure 6b). We could also detect higher MFIs of endogenous p65 in VP40-expressing cells (Figure 6c), a phenotype we corroborated in the context of trVLP infection (Figure S6) and quantitative image analyses (Fig. 6d). Based on our previous data (46), we hypothesized that CD81 might inhibit NFκB-activation imposed by EBOV VP40. To test for this, we activated NFκB by transfection of VP40 and co-transfected 293T to express increasing amounts of CD81 (Figure 6e). Of note, exogenous gradual CD81-expression reduced VP40-mediated NFκB-signaling. This effect was specific for VP40-mediated NFκB-activation, whereas CD81 did not suppress induction of NFκB triggered through constitutively active IKKβ (Figure 6e). Altogether, EBOV VP40 activates NFκB signaling in 293T cells, in a way that is negatively regulated by CD81.

### CD81 restricts early steps of EBOV trVLP infection

For some viruses, i.e. HCV and HPV, tetraspanins play important roles during viral entry and early infection (97–100). To explore whether CD81, CD63 or CD9 are involved in early steps of EBOV infection, including entry, EBOV trVLP-GFP infection in 293T control, CD81 KO, CD63 KO and CD9 KO cells was analyzed. In trVLP infection, the minigenome is replicated and transcribed only by pretransfected EBOV RNP and no T7/pol II mediated initial transcription is involved. Of note, KO of CD81, but not of CD63 and CD9, resulted in increased GP and VP40 expression in trVLP infected cells (Figure 7a, left and quantification below). Similarly, KO of CD81 led to increased trVLP-GFP infection rates (Figure 7a, right). An increased infection rate could result from more efficient entry and/or from altered minigenome replication/transcription, which we already demonstrated to be restricted by CD81 (Figures 3 and 6). To specifically assess EBOV GP-mediated entry, 293T control, CD81 KO, CD63 KO and CD9 KO cells were infected with GFP expressing lentiviruses (lenti-GFP) pseudotyped with either EBOV GP or VSV G. The infection rates were analyzed by flow cytometry. KO of CD81, but not of CD63 and CD9, enhanced EBOV GP-, but not VSV G-mediated entry of lenti-GFP (Figure 7b), suggesting a suppressing role of CD81 in EBOV GP-mediated entry. Vice versa, we employed trVLP-dGP-GFP pseudotyped with EBOV GP or VSV G for infection revealing that CD81 KO enhanced the infection of trVLP-dGP-GFP irrespective of the GP used (Figure 7c). Authentic EBOV or EBOV VLPs are internalized mainly through macropinocytosis (18, 19, 101). We therefore studied whether CD81 has an effect on macropinocytosis by analyzing the uptake of Dextran-555, a fluid phase marker, by flow cytometry. KO of CD81 enhanced the uptake of Dextran-555 in 293T cells (Figure 7d), suggesting a role of CD81 in macropinocytosis. Dextran-555 uptake in Huh7.5 cells was less dependent on CD81 (Fig. S7a) whereas in primary monocyte-derived macrophages (MDM) Dextran-555 internalization was clearly boosted upon CD81 KO (Fig. S7b). In addition, we analyzed if infection of 293T cells with EBOV GP pseudotyped trVLPs is reduced by CD81 overexpression (Figure S7c) and found, that high CD81 levels suppress trVLP infection in 293T cells (Figure 7e). This confirms the negative role of CD81 in EBOV trVLP infection in 293T cells. Altogether, the cumulated data indicates that CD81 restricts EBOV trVLP cellular uptake and entry.

**Fig. 7.**
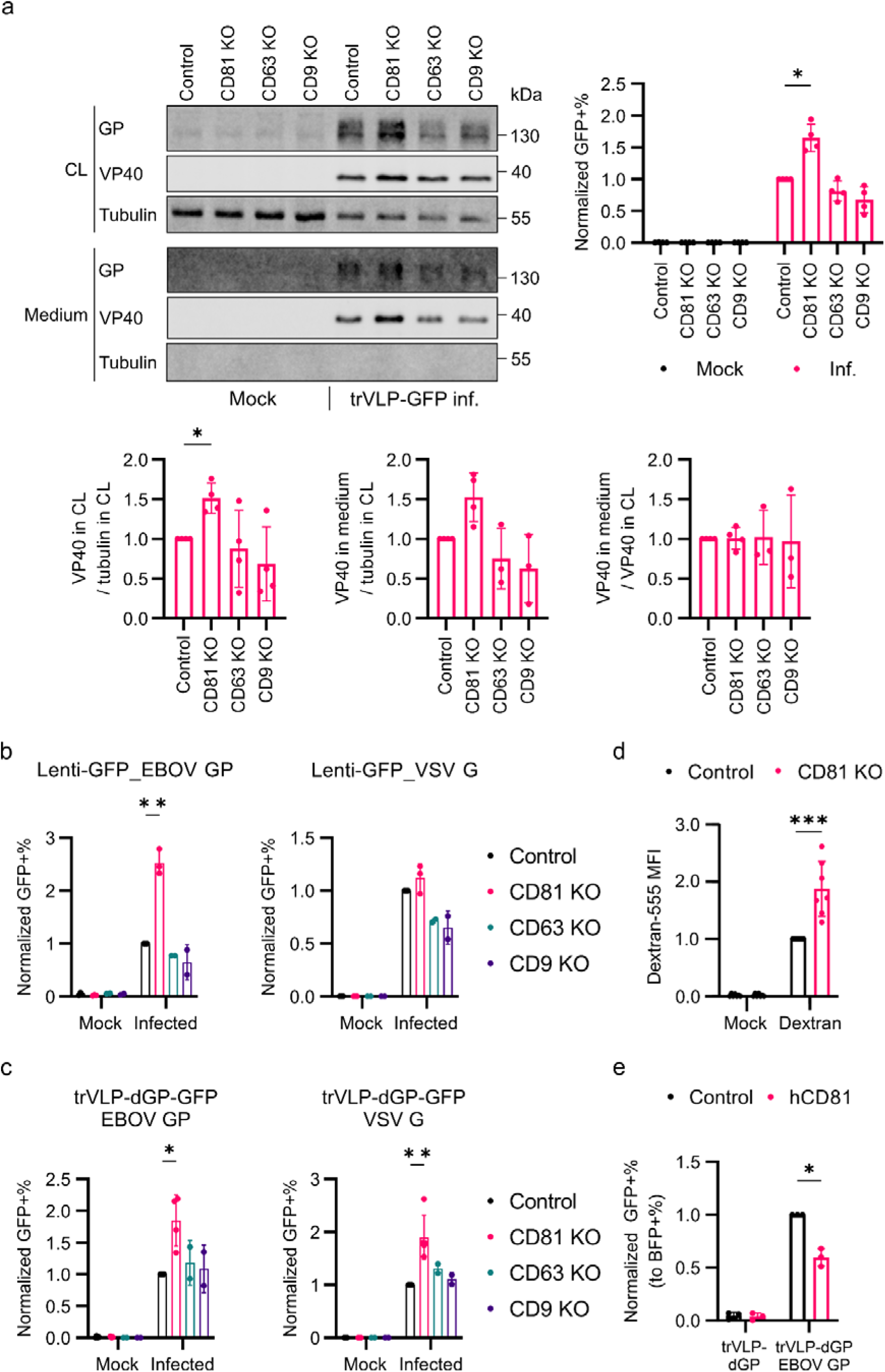
CD81 restricts early steps of EBOV trVLP infection. (a) 293T control, CD81 KO, CD63 KO and CD9 KO cells were mock or pre-transfected to express EBOV RNPs and Tim1 (EBOV attachment factor). 1 day later, the cells were infected with trVLP-GFP. 4 d.p.i., an aliquot of the cells was harvested for flow cytometry and the remaining cells and medium were harvested for WB. Indicated are representative WB images and mean values (normalized to inf. control cells) of WB signal quantification and infection rate +/- SD (n=4). RM one-way ANOVA was used for statistical analysis. (b) 293T control, CD81 KO, CD63 KO and CD9 KO cells were mock infected or infected with lenti-GFP pseudotyped with EBOV GP or VSV G. 3 d.p.i., the cells were harvested for flow cytometry. Indicated are mean values of infection rate (normalized to infected control cells) +/- SD (n=3 for 293T control/CD81 KO, n=2 for CD63/CD9 KO). (c) 293T control and KO cells were mock or pre-transfected to express EBOV RNP and Tim1. 1 day later, the cells were mock infected or infected with trVLP-dGP-GFP pseudotyped with EBOV GP or VSV G. 3 d.p.i., cells were harvested for flow cytometry.. Indicated are mean values of infection rate (normalized to infected control cells) +/- SD (n=4 for 293T control/CD81 KO, n=2 for CD63/CD9 KO). (d) 293T control and CD81 KO cells were left untreated (mock) or incubated with Dextran-555 (10K, 0,4 mg/ml). After incubation at 37 °C for 15 min, the cells were harvested for flow cytometry to analyze Dextran uptake. Indicated are mean values (normalized to control cells added with Dextran) of MFI +/- SD (n=7). (e) 293T WT cells were pre- transfected with plasmids expressing EBOV RNP, Tim1 and BFP and a CD81-HA expression plasmid or the control vector. 1 day later, cells were treated with trVLP-dGP-GFP or trVLP-dGP-GFP pseudotyped with EBOV GP. 48 h.p.i. cells were harvested for flow cytometry. Indicated are mean values of infection rate (normalized to BFP+% and further control infected cells) +/- SD (n=3). Welch’s t-test was used for statistical analysis: <0.05 (*), <0.01 (**), <0.001 (***).

### CD81 antibody treatment inhibits EBOV infection

Since CD81 itself restricts EBOV trVLP infection at early and late steps of viral replication, we hypothesized that targeting CD81 with a crosslinking antibody that induces cell signaling could exert antiviral effects. The antibody 5A6 was initially identified to bind to CD81 and induce antiproliferative effects (102). 5A6 was further shown to bind to the large extracellular loop of CD81 and induce conformational changes and CD81 clustering at the cell surface (103, 104). Another study stated that 5A6 activates a caspase 3-PARP-triggered pathway (105). Strikingly, CD81 5A6 pre-treatment at 1 or 5 µg/ml suppressed EBOV trVLP infection of CD81-expressing 293T cells when VLPs were pseudotyped with either GP or VSV G (Figure 8a - c). Importantly, the effect is CD81 specific, since an isotype control antibody did not exert any antiviral effects and the CD81 antibody did not reduce EBOV trVLP infection in CD81 KO cells. Even though, as expected, infection levels were generally higher in CD81 KO cells (Figure 8b and c). Furthermore, moving to authentic infection systems, EBOV infection of 293T cells was also suppressed by CD81 antibody 5A6 treatment (Figure 8d and e). In contrast, neither the CD81 antibody, nor the CD81 KO had any suppressive effect on MeV infection (Figure 8f). Altogether, CD81 interferes with EBOV trVLP infection at multiple steps throughout the viral life cycle and targeting CD81 via the cell signaling-inducing antibody 5A6 specifically inhibits trVLP as well as authentic EBOV infection.

**Fig. 8.**
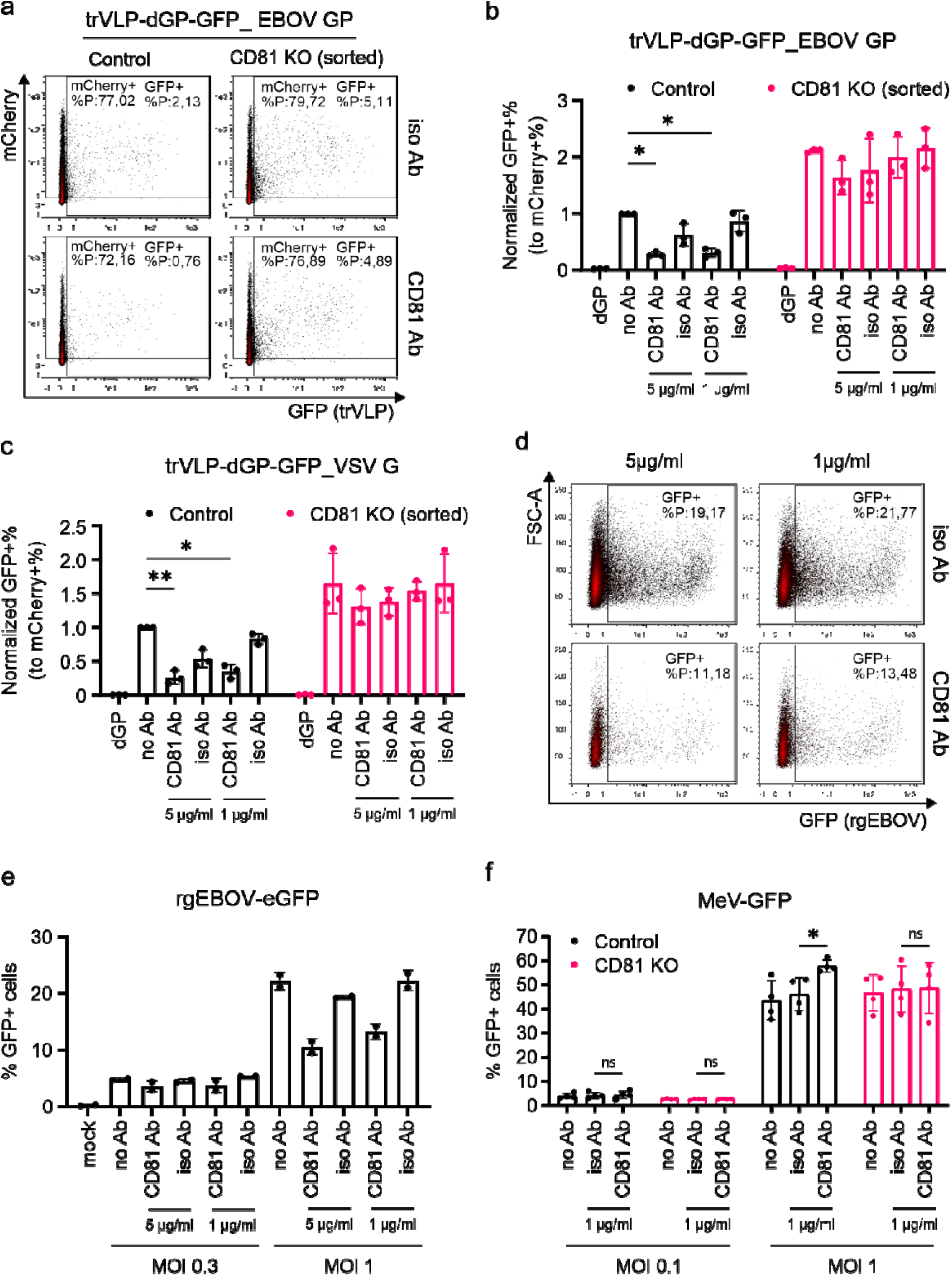
CD81 antibody treatment inhibits EBOV trVLP and authentic EBOV infection. (a-c) 293T control and CD81 KO cells were pre-transfected to express EBOV RNP, Tim1 and mCherry. 1 day later, the cells were mock treated or treated with the CD81 antibody (5A6) or the corresponding isotype antibody. After 1 h incubation, the cells were infected with trVLP-dGP-GFP or trVLP-dGP-GFP pseudotyped with EBOV GP (a,b) or VSV G (c). 2 d.p.i., cells were harvested for flow cytometry. Shown are (a) representative density plots and (b,c) mean values of infection rate (normalized to mCherry+ % and further No Ab treated control cells) +/- SD (n=3). (d,e) 293T cells were treated with no Ab, CD81 antibody (5A6) or the corresponding isotype antibody. 1 h later, the cells were mock infected or infected with rgEBOV-GFP at MOI 0.3 or 1. 1 d.p.i., cells were harvested and fixed for flow cytometry. Shown are (d) representative density plots and (e) mean values of infection rate (GFP+%) (n=2, in duplicates). (f) As specificity control, 293T control and CD81 KO cells were treated with no Ab, CD81 antibody (5A6) or the corresponding isotype antibody and infected 1 h later with MeV-GFP at MOI 0.1 or 1. 2 d.p.i., cells were harvested and fixed for flow cytometry. Shown are mean values of infection rate (GFP+%) +/- SD (n=4). (b,c,f) Two-way ANOVA with Dunnett correction was used for statistical analysis: <0.05 (*), <0.01 (**).

## Discussion

By comprehensive assessment of cell surface protein levels in EBOV GP expressing cells, we identified tetraspanins CD81, CD63 and CD9 to be dysregulated by EBOV GP. Not only EBOV GP but also MARV, SUDV, TAFV and RESTV GP were able to interfere with surface CD81, CD63 and CD9. This effect was not observed with glycoproteins from other virus families tested, suggesting a conserved function among filovirus GP. Of note, GP seems to exert specificity in lowering levels of cell surface receptors, as of 332 receptors assessed by flow cytometry, 46 showed more than 2-fold reduced antibody binding and three receptors were increased upon GP expression (Figure 1A, sup. Table 1). This is remarkable, as it was hypothesized that EBOV GP might form a glycan shield at the PM, that generally interferes with antibody binding and for instance lowers the accessibility of HLA towards cytotoxic T lymphocytes (CTL) (56, 60). Our study indicates that EBOV GP alters the composition of the PM by additional mechanisms. As exemplified for CD81, such mechanisms include, in addition to glycan shielding, direct or indirect protein interaction and therefore potentially interference with receptor localization, recycling and trafficking. The latter could involve GP-mediated alteration of protein turnover via the proteasomal and lysosomal routes. CD81 is not only modulated by GP, but also by VP40, indicating that Ebola exploit various independent mechanisms to antagonize and inactivate host cell suppression. It will be interesting to clarify the exact mechanistic interplay of these various modalities in the process of GP and VP40-mediated alteration of CD81. Such an approach should include extensive and systematic mutagenesis of GP, VP40 and CD81 to identify structural motifs that mediate CD81-modulation and their relative contribution to the mechanisms discussed above.

CD81 and CD9 share around 60% sequence similarity and both tetraspanins have similar structures, while especially the extracellular loop sequence differs from each other (106, 107). Therefore, since both CD81 and CD9 are downregulated by GP, even though only CD81 is strongly modulated in primary macrophages (Fig. 1d and f), it is tempting to hypothesize that the shared intracellular sequence motif (108) and transmembrane domain are involved in receptor modulation, while the unique sequence identity in CD81 (mainly located to the extracellular loops) confers the inhibitory role of CD81 on EBOV trVLP replication. In GP, mutations in the RBD, FP and TMD impaired even though did not disrupt CD81 modulation Figure S2e). Hence, multiple sequence motifs in GP seem to participate in the complex interaction with CD81, that conceivably will involve domains necessary for functional interaction (for instance GP2), while others might direct CD81 for proteasomal and lysosomal degradation and contribute to glycan shielding (GP1).

Our studies firmly establish CD81, but not CD63 or CD9 as inhibitory factor acting at multiple steps of the EBOV life cycle. Importantly, our findings are not at odds with a recent report suggesting a beneficial role of CD9 in acting as co-factor in VP40 VLP release (109). However, as CD9 KO in 293T cells did not result in pronounced phenotype in 293T or Huh7.5 cells (Fig. 3a and b), we focused on CD81. CD81 suppresses levels of EBOV RNA species, mainly the cRNA, while IB formation that is essential for efficient EBOV genome replication and transcription seems not affected by CD81 (31–33, 90, 93). Another restriction imposed by CD81 include GP-dependent and independent routes of entry, involving a not-yet discovered role of CD81 in macropinocytosis. In line with this, CD81 interacts with the GTPase Rac and regulates Rac activation, which is known to play an important role in macropinocytosis (110, 111). Besides, CD81 is also related to ERMs (Ezrin-Radixin-Moesin), actin binding proteins that are also involved in macropinocytosis (112, 113). More direct roles of CD81 in GP-mediated attachment and entry could be related to the CD81 interaction partners integrin α1, 4, 5, 6 and β1 (114–116). Integrin α5β1 is involved in EBOV entry by regulating cathepsins (56, 58, 117). Our flow cytometry screen includes antibodies against integrins, αβ1, β5 and β7 (Table S1). However, none of them is modulated by GP. Extending our analysis to other CD81 interaction partners that are non-TSPs and assessed by our approach, neither CD19 (118) nor CD44 (119) are GP-modulated (Table S1). This, and the fact that GP directly interacts with CD81 (Fig. 5), argues against a mechanism in which CD81-intereference is a bystander effect of GP modulating another CD81-interaction partner.

Of note, the multilayered inhibitory effects of CD81 on EBOV replication could be related to its ability to suppress NFκB signaling (46). NFκB is a pleiotropic transcription factor that has profound effects on cellular proliferation, apoptosis, organization of the cytoskeleton and is also affected by the endocytic machinery (120–123). On top, it is an important factor boosting the interferon antiviral response (124). In this regard, given that EBOV efficiently blunts the innate immune response (125–128), the sustained NFκB activation in the absence of innate immune signaling likely results in an overall proviral and replication supportive cellular environment (129, 130). 293T cells employed here neither express tetherin (also known as BST2) nor TLR4, that were previously implied to activate NFκB in cooperation with GP and VP40 (131, 132). We hence demonstrate that VP40 activates NFκB and this phenotype is confirmed in EBOV trVLP infected cells. Most importantly, this finding is supported by a recent study that extended this phenotype to VP40 proteins of other members of the *Filoviridae* and suggested a mechanism that is triggered via the TNFR1 signaling axis (133). Altogether, supported by the new data from the Ebihara lab (133), and given the importance of NFkB activation during EBOV infection in driving the cytokine storm and immunopathogenesis (134, 135), these findings reveal a previously unrecognized, TLR4-independent mechanism of N-kB activation by EBOV VP40.

Finally, it is striking that the cell signaling-inducing CD81 antibody 5A6, targeting the extracellular loop of CD81, suppressed infection of 293T cells with EBOV trVLPs pseudotyped with GP or VSV G, as well as authentic EBOV infection (Fig. 8). In contrast, pre-treatment with the 5A6 antibody had no effect on MeV replication, indicating a filovirus-specific activity. 5A6 has been shown to inhibit breast cancer cell invasion and migration, and B cell lymphoma growth with good safety profiles in mice via activation of cell signaling (104, 105). The functional relation of these findings to the restricting role of CD81 in the EBOV life cycle is not clear yet. Nevertheless, the data indicates that EBOV interferes with cell surface CD81 to avoid associated signaling which results in an inhibitory activity on the various steps of viral replication, involving NFκB. Findings that warrant further investigation. Our study has certain limitations. First, the exact structural and functional motifs in GP and CD81 that are involved in modulation and functional restriction remain largely elusive. Second, the restricting activity of CD81 on the EBOV life cycle was formally only demonstrated in the EBOV trVLP surrogate system of viral replication. Difficulties to assess the CD81 relevance in the context of authentic EBOV infection are due to the high potency of CD81-modulation by GP and VP40, which makes it challenging to draw conclusions when comparing EBOV infection in control vs CD81 KO cells. Nevertheless, a strong argument for CD81 also restricting authentic EBOV infection is the fact that we measure robust downmodulation of CD81 in the context of authentic EBOV infection not only in 293T cells but also primary human macrophages (Fig. 1d and f) and inhibition of authentic EBOV infection specifically by the CD81-targeting antibody (Fig. 8d and e). A finding that has certain translational potential that will be further followed up.

In summary, we identified novel EBOV GP modulated plasma membrane proteins. Three of them are the tetraspanins CD81, CD63 and CD9 that are downregulated by highly divergent filovirus GPs. CD81, and not CD63 nor CD9, emerges as cellular factor with inhibitory activity at multiple steps of the EBOV life cycle that are interconnected to NFκB and therefore filoviral immunopathogenesis.

## Consent for publication

All authors gave their consent to publish. All authors read and approved the final manuscript.

## Availability of data and material

All data generated and analyzed during this study are included in this published manuscript.

## Authorship inclusion and ethics statement

DH and EH performed most of the experiments supported by MBo and MBu. JB, LWi, LWe and TH did the authentic EBOV infection experiments. JL did the initial screen to find EBOV GP modulated receptors. JK and LWi analyzed TSP versus GP interaction and supported mechanistic studies. DH, EH, LWe, TH and MS planned the experiments and analyzed the data. MS wrote the manuscript draft. MS supervised the overall study. TH and MS provided resources. All authors contributed to editing and developed the manuscript to its final form.

## Acknowledgements

We thank Daniel Sauter (Tübingen), Thomas Gramberg (Erlangen) and Stefan Pöhlmann (Göttingen) for the kind contribution of cell lines, reagents and plasmids. Furthermore Ulrich Lauer (Tübingen) for support with MeV-GFP.

## Figures and Legends

**Sup. Fig. S1.**
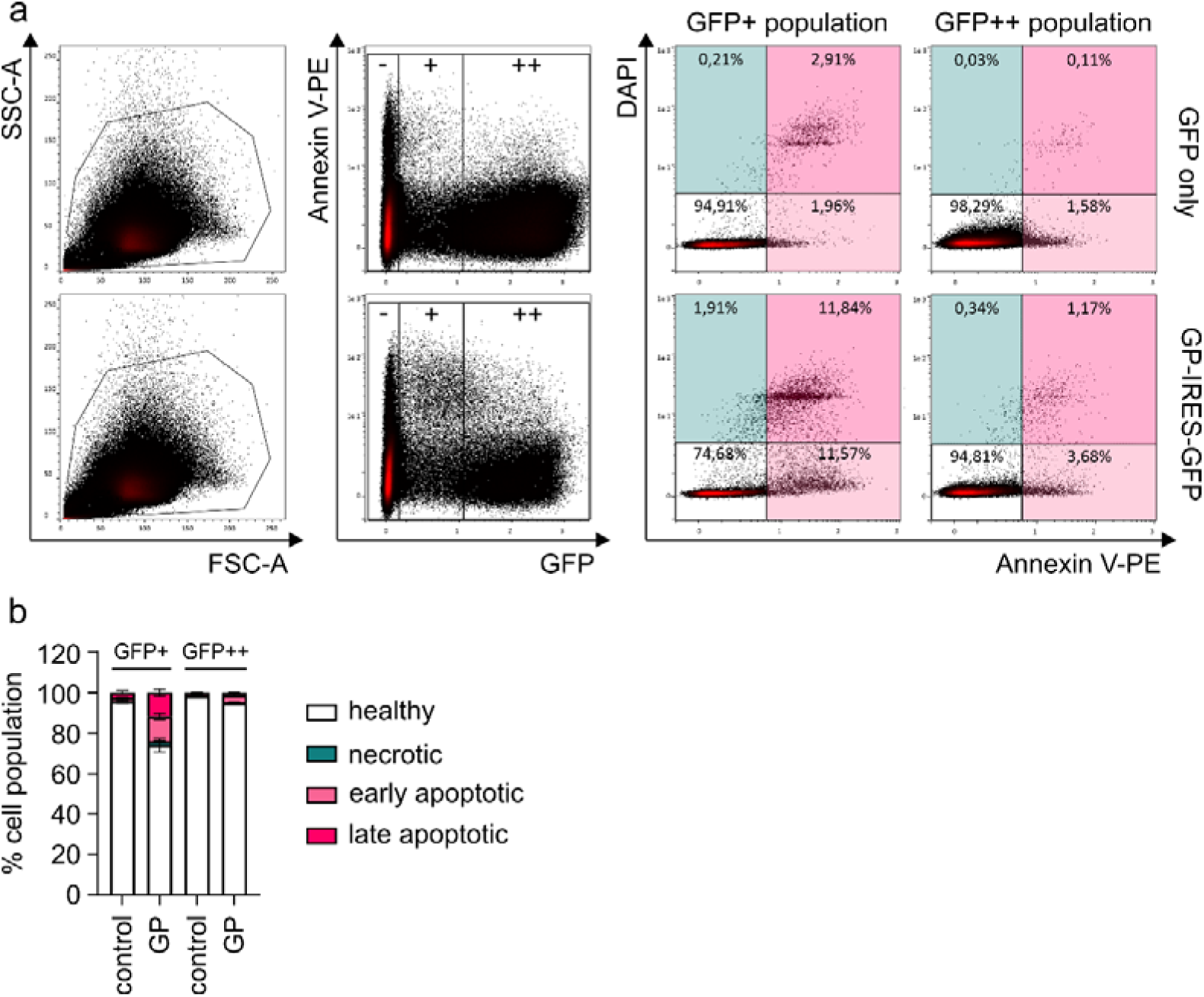
Flow cytometry gating and GP-induced cytotoxicity. (a) 293T cells were GFP only transfected (top) or transfected to co-express GFP and EBOV GP (bottom). The general population of living cells was gated according to SSC-A and FSC-A (left panel of FACS plots). 2 d.p.t., the cells were stained with Annexin V (AnV)-PE and DAPI to detect healthy (AnV-/DAPI-), necrotic (AnV-/DAPI+), early apoptotic (AnV+/DAPI-) or late apoptotic (AnV+/DAPI+) cells (right panel of FACS plots). We further included two regions of medium (+) or high (++) levels of GFP, hence GP-expression. (b) Quantification of the percentage of healthy, necrotic, early and late apoptotic cells in GFP only control or GP-expressing 293T cells according to regions of medium (GFP+) or high (GFP+) levels of GFP/GP expression. The diagram shows stacked columns indicating the relative percentage of the respective cell fraction +/- SD of three independent experiments.

**Sup. Fig. S2.**
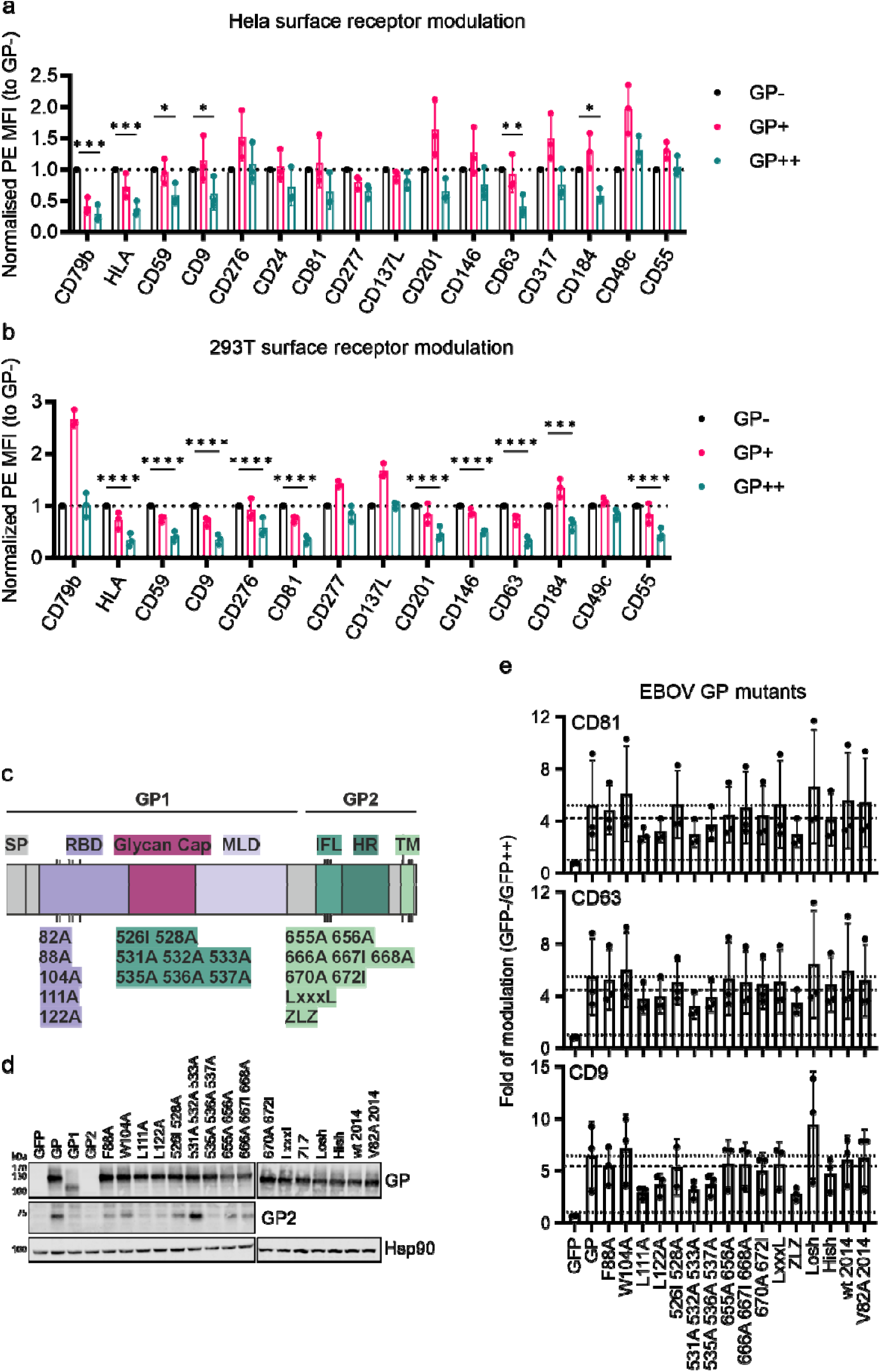
Modulation of surface receptors by EBOV GP and GP mutants. (a,b) Relative expression (PE MFI normalized to GP- cells) of surface receptors (2-fold higher than unstained ctrl) in (a) Hela (b) and 293T cells according to no (-), medium (+) and high (++) levels of GFP and hence GP expression. Shown are mean values +/- SD (n=3). Two-way ANOVA with Dunnett correction was used for statistical analysis: <0.05 (*), <0.01 (**), <0.001 (***), <0.0001 (****). (c) Scheme of EBOV GP domain structure with indicated mutations. Structural domains are highlighted in color: signal peptide (SP), receptor-binding-domain (RBD), mucin-like domain (MLD), internal fusion loop (IFL), heptad repeats (HR) and transmembrane domain (TM). (d) 293T cells were transfected to express indicated GP subunits and mutants. 24 h.p.t. cells were harvested for WB analysis. Shown are representative WB images (n=2). (e) Downregulation of surface CD81, CD63 and CD9 by EBOV GP mutants. Shown are mean values of fold of modulation (GFP-/GFP++) +/- SD (n=3).

**Sup. Fig. S3.**
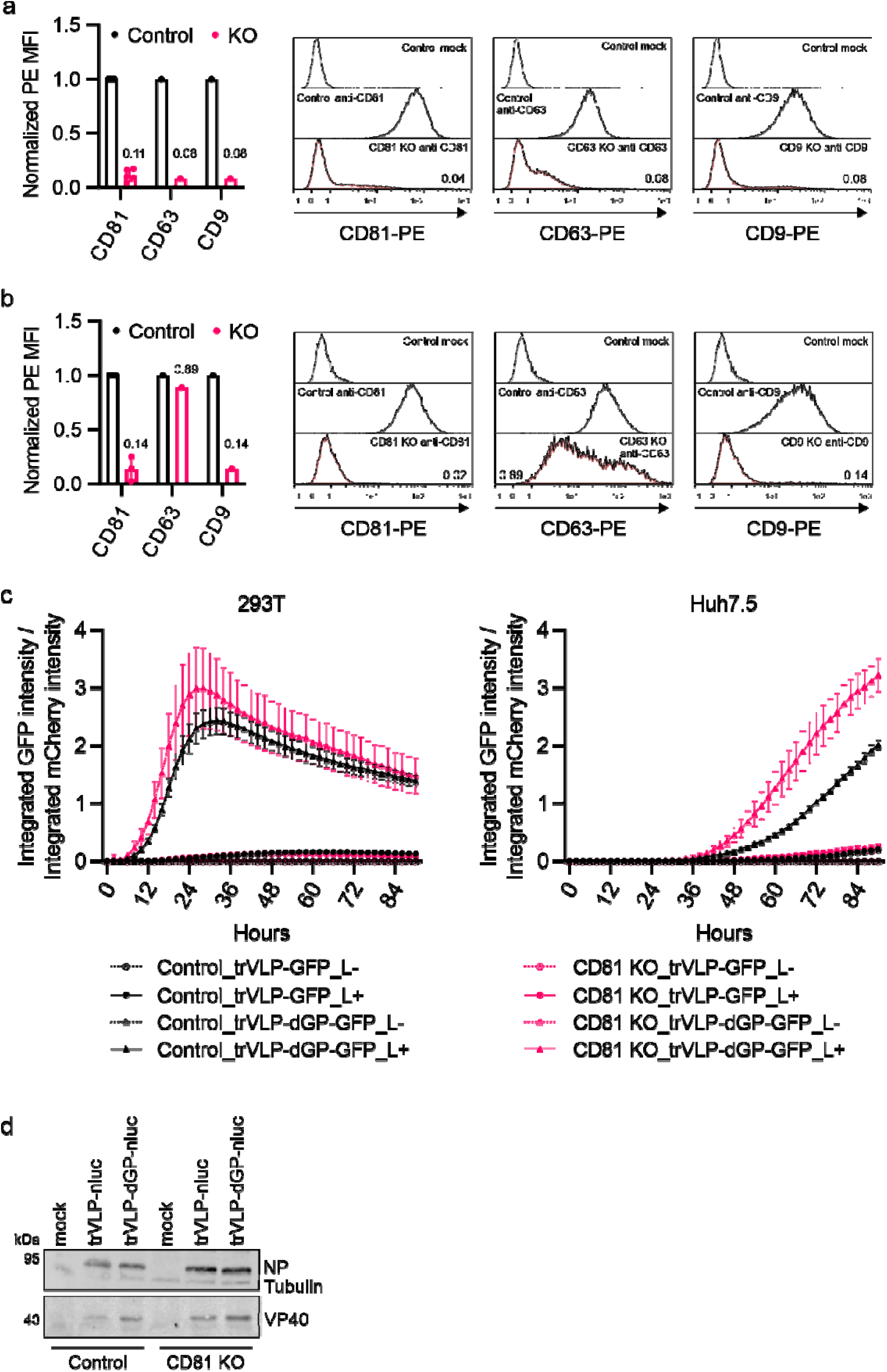
The role of CD81 in EBOV trVLP replication. (a) To verify the KO-efficiency, 293T (n=5 for control/CD81-KO, n=1 for CD63/CD9-KO) and (b) Huh7.5 cells (n=3 for control/CD81-KO, n=1 for CD63/CD9-KO) were stained with PE conjugated antibodies against CD81, CD63 and CD9 for flow cytometry analysis. Shown is PE MFI (normalized to control cells) or mean values of normalized PE MFI +/- SD and histograms. (c) 293T (n=4) and Huh7.5 (n=3) control and CD81 KO cells were transfected with plasmids for trVLP-GFP and trVLP-dGP-GFP replication and mCherry expression plasmid (transfection ctrl). Transfection without viral RNA polymerase L (L-) was included as negative ctrl. 6 h.p.t., the cells were imaged every two hours. Integrated green intensity (GFP expression) normalized to integrated red intensity (mCherry expression) was plotted, shown are mean values +/-SEM. (d) 293T control and CD81 KO cells were transfected for trVLP-nluc and trVLP-dGP-nluc replication. 48 h.p.t., cells were harvested for WB analysis. Shown are representative WB images (n=3).

**Sup. Fig. S4.**
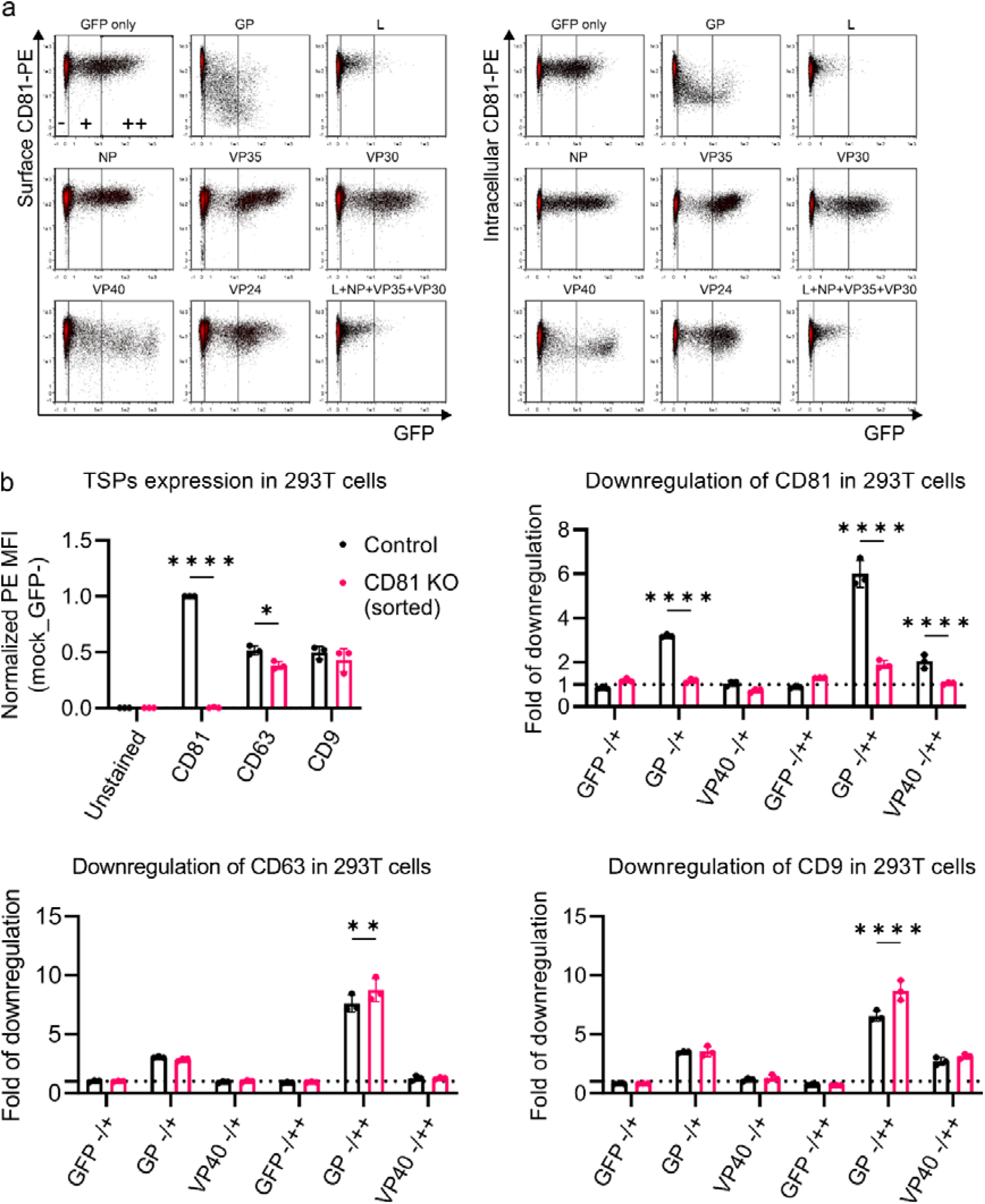
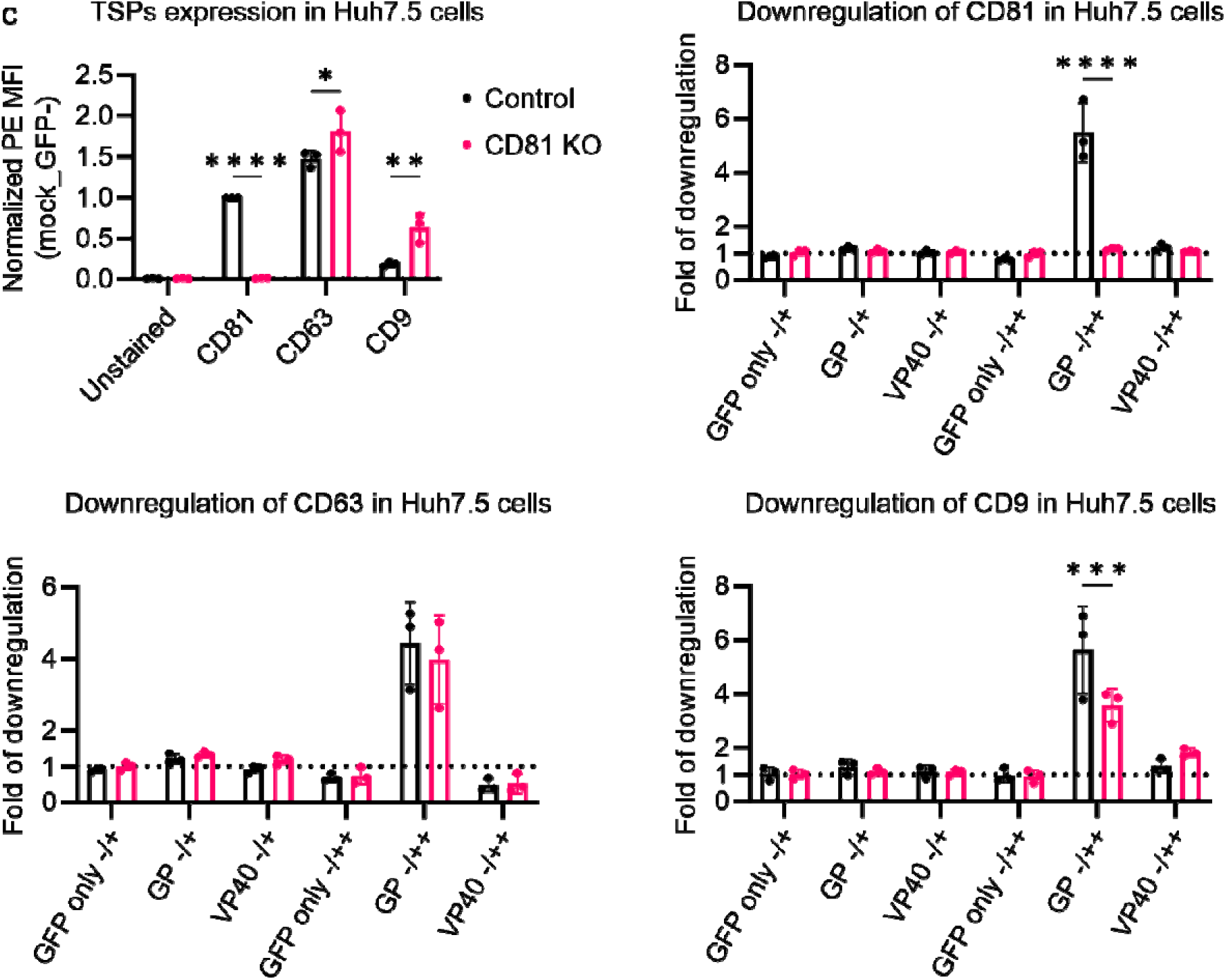
Downregulation of CD63 and CD9 is not dependent on CD81. (a) Modulation of surface and intracellular CD81 by EBOV proteins in 293T cells, shown are representative images of flow cytometry density plots. (b) 293T and (c) Huh7.5 cells were mock transfected or transfected to express GFP only or co-express GFP and EBOV GP or VP40. 2 d.p.t., the cells were harvested and stained for CD81, CD63 and CD9 for flow cytometry analysis. Shown are mean values of normalized PE MFI (to CD81-PE MFI of mock transfected control cells (GFP-)) and fold of modulation (in PE MFI) +/- SD (n=3). Two-way ANOVA with Šidák correction was used for statistical analysis: <0.05 (*), <0.01 (**), <0.001 (***), <0.0001 (****).

**Sup. Fig. S5.**
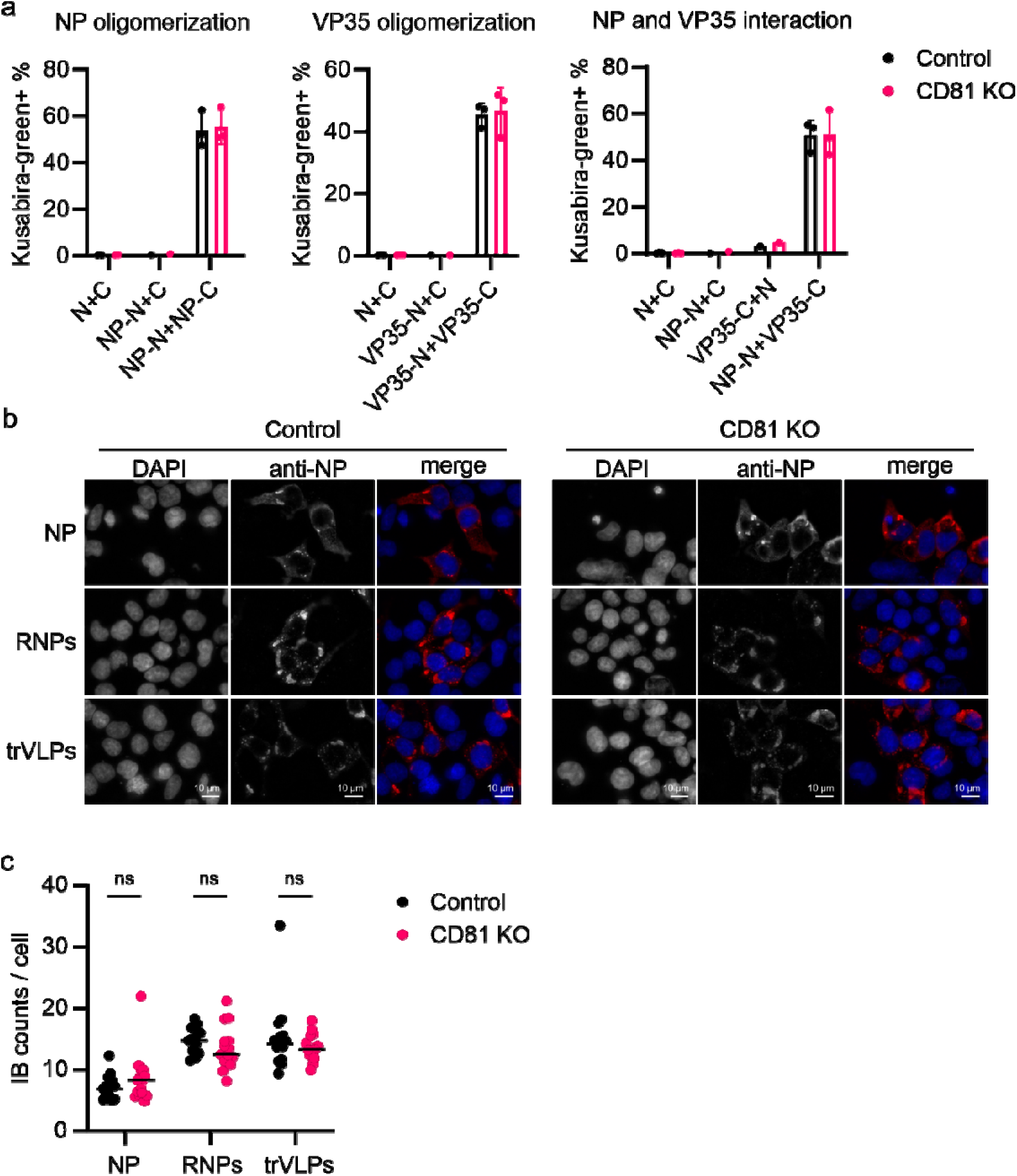
The effect of CD81 on NP and VP35 interaction and inclusion body (IB) formation. (a) 293T control and CD81 KO cells were transfected to express the N- and C-part of Kusabira-green (KG) fused to N or VP35 as indicated for the BiFC assay. 2 d.p.t., the cells were harvested for flow cytometry analysis. Shown are % of KG-positive cells +/- SD (n=3). (b) 293T control and CD81 KO cells were transfected to express NP alone, all ribonucleoprotein (RNP) components (NP, VP35, VP30 and L), or transfected with RNPs and subsequently infected with trVLPs. Cells were fixed 24 h.p.t. and stained for NP and DAPI. Shown are representative images (n=3). (c) The number of IBs per cell was quantified using ImageJ. Two-way ANOVA with Šidák’s correction was used for statistical analysis.

**Sup. Fig. S6.**
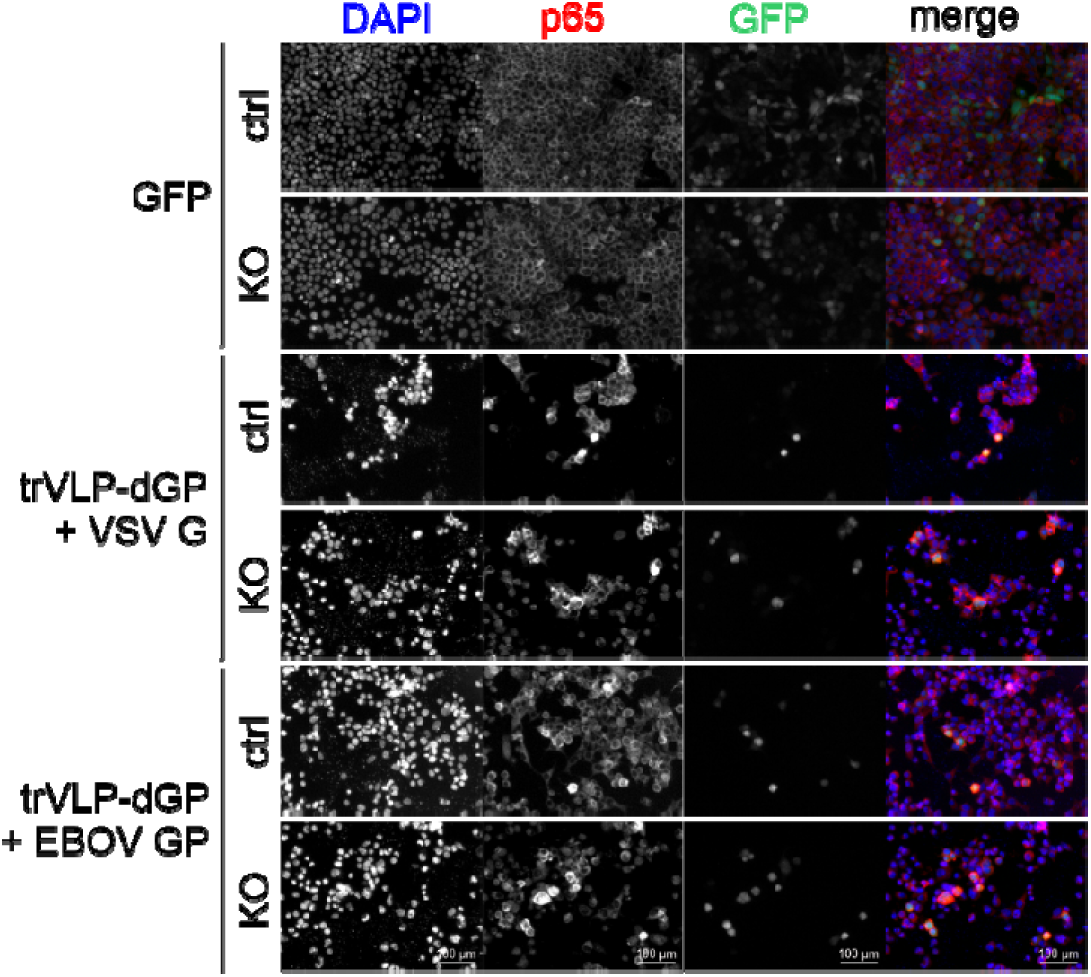
Induction of p65 by trVLP infection. 293T control and CD81 KO cells were transfected to express GFP as a control or with RNP components (NP, VP35, VP30, L) and subsequently infected with trVLPs pseudotyped with either VSV G or EBOV GP. 48 h.p.i. cells were fixed, stained for p65 and DAPI and imaged by fluorescence microscopy. For quantification of IF signal see Figure 4D. Shown are representative images (n=3).

**Sup. Fig. S7.**
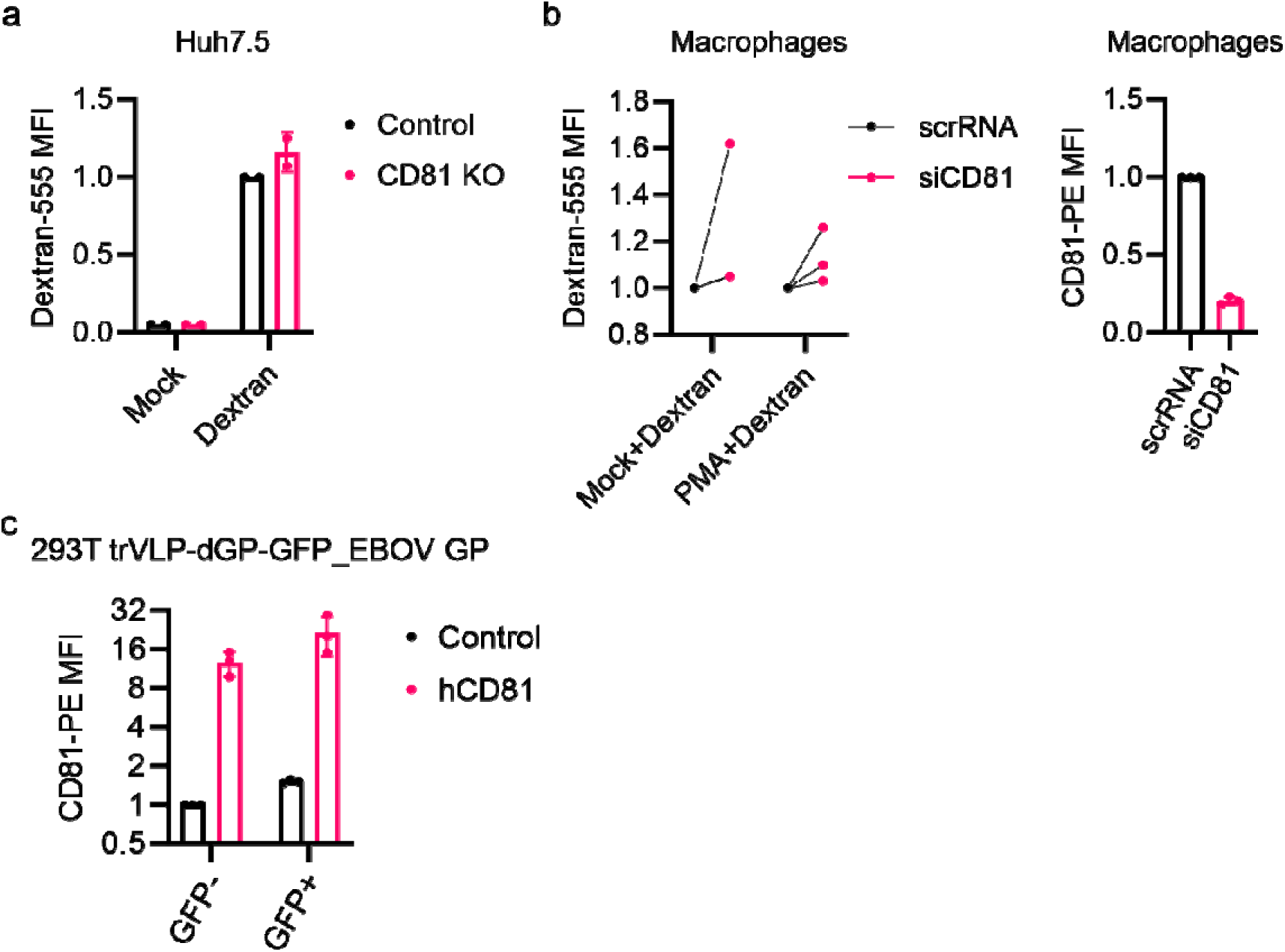
The role of CD81 in EBOV trVLP infection. (a) Huh7.5 control and CD81 KO cells were left untreated or incubated with Dextran-555 (10K, 0,4 mg/ml). After incubation at 37 °C for 15 min, the cells were harvested for flow cytometry to analyze Dextran uptake. Shown are MFI values (normalized to control cells added with Dextran) +/- SD (n=2). (b) Human primary macrophages were transfected with non-targeting siRNA (scrRNA) or siRNA targeting CD81 (siCD81). 4 d.p.t., the cells were mock treated or treated with PMA (1 μM). After 30 min incubation, the PMA was removed and Dextran-555 was added. 15 min later, the cells were harvested for flow cytometry to analyze Dextran uptake. Shown is Dextran-555 MFI (normalized to scrRNA transfected cells) of macrophages from three independent donors (left). Macrophages without PMA and Dextran-555 treatment were harvested and stained for CD81 for flow cytometry. Shown is CD81-PE MFI (normalized to scrRNA) of macrophages from the three donors. (c) CD81 expression of 293T cells, pre-transfected with empty vector (Empty) or human CD81 (hCD81) expression plasmid and infected with trVLP pseudotypes, was analyzed by flow cytometry. Shown are mean values of CD81-PE MFI (normalized to control non-infected (GFP-) cells) +/- SD (n=3).

